# Description of canine- and feline-derived strains of the bile acid-converting bacterium *Peptacetobacter hiranonis: P. hiranonis* subsp. *deconjugans* subsp. nov. and *P. hiranonis* subsp. *nondeconjugans* subsp. nov.

**DOI:** 10.64898/2026.08.21.746369

**Authors:** Bruna Correa Lopes, Jonathan Turck, Amanda Blake, Luis Fernando da Costa Medina, Sara D. Lawhon, Jan S. Suchodolski, Rachel Pilla

## Abstract

The bile acid-converting *Peptacetobacter hiranonis* is a Gram-positive, anaerobic, potentially spore-forming bacterium. It was first isolated from human feces and was subsequently shown to convert bile acids (BA) in both *in vitro* and *in vivo* experiments. The conversion of BA relies on the presence of the 7α-dehydroxylation multi-step pathway, encoded by the BA-inducible (bai) operon, harbored by *P. hiranonis*. In companion animals, *P. hiranonis* has been characterized as a biomarker for intestinal health, with its loss associated with dysbiosis. However, characterization of *P. hiranonis* cultured from companion animals is limited. An in-depth characterization of *P. hiranonis* was published by Chen *et al.* recently, including the proposal of a new species, *Peptacetobacter hominis*. We have sequenced the whole genome of both canine- and feline-derived strains of *P. hiranonis*, characterized these strains biochemically, and assessed their *in vitro* BA-converting ability as well as their antimicrobial resistance profiles. The strains described here can convert primary into secondary BAs and are whole-genome inhibited by low concentrations of amoxicillin-clavulanate, cefepime, ceftriaxone, chloramphenicol, ciprofloxacin, clindamycin, and metronidazole. Based on whole genome analysis, we propose dividing *P. hiranonis* into two host-adapted subspecies: *P. hiranonis* subsp. *deconjugans* and *P. hiranonis* subsp. *nondeconjugans*, based on their genomic differences and divergent ability to deconjugate BAs – a function that appears widely distributed among *P. hiranonis* strains cultured from dogs, but absent from those cultured from cats. Taken together, our results confirmed the BA conversion ability of *P. hiranonis* cultured from dogs and cats and reveal host-associated genomic and functional differences within the species.

## INTRODUCTION

The intestinal microbiota of dogs and cats harbors various microbial species involved in maintaining intestinal balance, including the bile acid (BA)-converting species *Peptacetobacter hiranonis*. *P. hiranonis*, previously known as *Clostridium hiranonis*, is an anaerobic, potentially spore-forming, and gram-positive bacterium [1,2]. In dogs and cats, *P. hiranonis* is described as a biomarker for gastrointestinal functionality and is closely linked to maintaining a balanced gastrointestinal microbiota [3].

*P. hiranonis* occupies a pivotal role in the conversion of primary [cholic acid (CA) and chenodeoxycholic acid (CDCA)] into secondary [deoxycholic acid (DCA) and lithocholic acid (LCA), respectively] fecal unconjugated BAs, and it is described as the main species with this ability within the canine and feline intestinal microbiome [4–9]. The conversion of BAs relies on the presence of the BA-inducible (bai) operon, which encodes a multi-step enzymatic pathway, and it is carried by *P. hiranonis* strains. In addition to its ability to convert primary into secondary unconjugated BAs, some strains of *P. hiranonis* can also deconjugate glycine- and taurine-conjugated BAs [2,10–12].

The lack or reduction of *P. hiranonis* and dysmetabolism of BAs have been associated with chronic inflammatory enteropathies in companion animals [13], and dysmetabolism of BAs has been linked to other chronic inflammatory diseases, including obesity, type 2 diabetes, and liver diseases in humans [14,15]. The disruption of microbial BA metabolism has been associated with reduced *P. hiranonis* abundance in several studies [8,13] and is commonly observed after antimicrobial use or chronic inflammatory enteropathies in dogs and cats [5,6,16].

Species within the genus *Clostridium* have been reclassified under different taxonomy recently; for example, *Clostridioides difficile* and *Peptacetobacter hiranonis* were previously known as *Clostridium difficile* and *Clostridium hiranonis*, respectively. The mentioned changes in *P. hiranonis* taxonomic classification refer to the creation of a new family called Peptostreptococcaceae under the Eubacteriales order, Clostridia class, and Firmicutes phylum, based on a culture-independent method, *i.e.,* genomic characterization [17]. Members of the Peptostreptococcaceae family were previously classified as *Clostridium* cluster XI.

The delineation of bacterial species usually relies on the average nucleotide identity (ANI) and digital DNA-DNA hybridization (dDDH), with proposed species boundary values of 95% for ANI [18–20] and 70% for dDDH [21,22]. These values may differ depending on the target bacterial species used for species or subspecies delineation analysis. Boundary values for subspecies classification have not yet been established. At the time of writing, the Peptostreptococcaceae family contains 19 different genera; among them, *Peptacetobacter,* which comprises *Peptacetobacter hominis* and *P. hiranonis* species.

*P. hiranonis* was first isolated from human feces in 1981 by Hirano *et al*. as bacteria capable of converting BAs [10]. Later, Kitahara *et al.* reported that although *P. hiranonis* can be detected in humans, it is usually present in smaller amounts and has a relatively low prevalence [23]. In companion animals, *P. hiranonis* is considered a core member of the healthy gut microbiome in both dogs and cats [24,25]. *P. hiranonis* has been described as a biomarker for intestinal functionality [3], and the restoration of *P. hiranonis* in dogs with chronic enteropathy has been linked to a better clinical response to fecal microbiota transplantation in these animals [26].

Despite its importance in canine and feline gut health, the biochemical and genomic characterization of *P. hiranonis* strains cultured from dog and cat feces is poorly described in the literature. Our objective in this study is (1) to compare genomes from *P. hiranonis* strains cultured from dogs and cats to the type strain and between them, (2) to describe the biochemical profile of *P. hiranonis* strains cultured from dogs and cats, (3) to assess the deconjugation and conversion ability of *P. hiranonis* strains cultured from dogs and cats, and (4) to assess the antimicrobial susceptibility profile of *P. hiranonis* strains cultured from dogs and cats.

## MATERIALS AND METHODS

### *Peptacetobacter hiranonis* Culture and Identification

Culture and isolation of *P. hiranonis* strains from the feces of dogs (*n =* 11) and cats (*n =* 11; Supplementary Table 1) was performed using conventional bacterial culture. Fecal samples were serially 10-fold diluted in phosphate-buffered saline (0.008 M sodium phosphate, 0.002 M potassium phosphate, 0.14 M sodium chloride, 0.01 M potassium chloride, pH 7.4, 500 ml; Pierce^TM^, IL) and plated onto *Brucella* blood agar (BBA) plates (Anaerobic Systems, CA; Supplementary File 1).

To identify *P. hiranonis*, colonies morphologically compatible with *P. hiranonis* were isolated and collected simultaneously for further identification using molecular methods. DNA extraction from each colony was performed as described by Dashti *et al.* [27]. Confirmation of *P. hiranonis’* 16S ribosomal RNA, *P. hiranonis’* bile salt hydrolase (BSH) gene, and a gene of the *P. hiranonis’* BA-inducible operon (*baiCD*) by qPCR was conducted following the protocol described by AlShawaqfeh *et al.* [28] and Correa Lopes *et al.* [9] (Table 1).

**Table 1.**
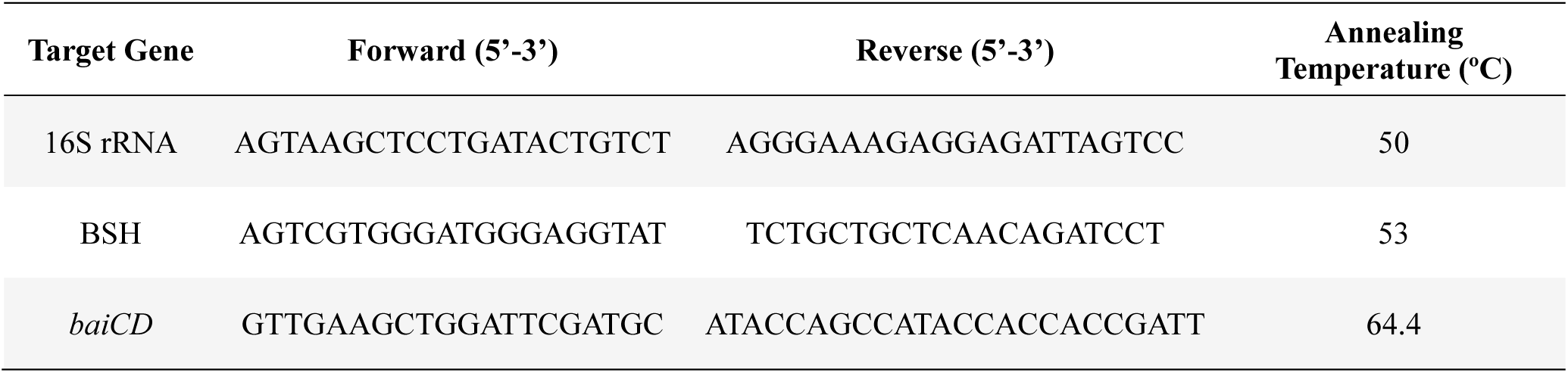
Forward and reverse primers were used to confirm the identity *of Peptacetobacter hiranonis* isolates cultured from the feces of dogs and cats.

The qPCR assay was performed using SsoFast EvaGreen^®^ Supermix (Bio-Rad Laboratories, CA) as described by AlShawaqfeh et al. [28,29]. For the qPCR reaction, 2 μl normalized DNA (5 ng/μl) was added to 5 μl SsoFast EvaGreen Supermix (Bio-Rad Laboratories), along with 0.4 μl each of forward and reverse primers (400 nM) as described in Table 1, and 2.2 μl DNA-free water. The Bio-Rad CFX384 Touch Thermal Cycler (Bio-Rad Laboratories) was used in this study. The cycling protocol is described as follows: initial denaturation at 98°C for 2 minutes; 35 cycles with denaturation at 98°C for 3 seconds; and annealing for 3 seconds.

### Sequencing *Peptacetobacter hiranonis* Genomes Using PacBio Sequel II

An aliquot of each strain of *P. hiranonis* (*n* = 21), 50 μL, was centrifuged at 5000g for 5 minutes. The supernatant was discarded, and the pellet was resuspended in 160 μL P1 buffer (QIAGEN, Germany). High molecular weight DNA was extracted using MagAttract HMW DNA Kit (QIAGEN). The DNA was eluted in 100 μL AE buffer (QIAGEN). DNA concentration was determined using the Invitrogen^TM^ Qubit^TM^ dsDNA HS Assay Kit (Life Technologies, USA).

To determine the size of the DNA, the sample was run on E-Gel SizeSelect 2% Agarose Gel (Invitrogen^TM^) along with a 1Kb ladder. The samples showed a DNA band higher than 15000 bp and were sheared using the Covaris G-tube (Covaris^®^, USA). The average size of the sheared DNA sample was determined using an Agilent 2100 Bioanalyzer (Agilent Technologies, USA). 1500 ng of the sheared and purified DNA was used as input for the library preparation using SMRTbell Express Template Prep Kit 2.0 (Pacific Biosciences, USA).

During library preparation, samples underwent DNA damage and end repair as well as barcode adapter ligation. To remove the unligated DNA fragments from the library, nuclease treatment was performed. The SMRT library was size-selected (> 6 kb) using a 0.75% Agarose gel in Blue Pippin (Sage Science). Following the size selection, the concentration of the final library was measured using the Qubit® dsDNA HS Assay Kit (Life Technologies). The library was then sequenced using the 30-hour movie time on the PacBio Sequel II (Pacific Biosciences).

### Assembly of *Peptacetobacter hiranonis* Genomes Using Flye

The method employed in the sequencing of *P. hiranonis* allowed coverage of base-called reads superior to 1000x coverage. The assembler Flye version 2.9.5 [30] for long reads was employed to generate consensus contigs with “--pacbio-hifi” parameter to attempt to assemble the genome from a file containing PacBio Hifi reads and “--genome-size 2.8m” set as the estimated genome size of *P. hiranonis* based on *P. hiranonis* DSM 13275 (cultured from human feces) and DGF055142 (cultured from dog feces). The “--asm-coverage” from 100 to 200 parameter was used to reduce coverage for initial disjoint assembly.

In summary, Flye generates a draft genome that uses only solid k-mers, a consensus process that will allow the selection of reads that belong to a defined contig. By default, Flye also runs one polishing iteration as the final assembly stage.

### Comprehensive Genome Analysis of *Peptacetobacter hiranonis* Using BV-BRC

Assembled contigs were used for the Comprehensive Genome Analysis on the Bacterial and Viral Bioinformatics Resource Center (BV-BRC) [31]. The RAST toolkit was used to identify repeat regions [32–34]. After repeat regions were identified, coding sequences (CDS) were called following Prodigal and Glimmer [35,36]. Antimicrobial resistance was projected by the Adaboost machine-learning [37], followed by an initial protein annotation using BLAT [38] and BLASTP [39] to identify CDSs that have homology to proteins in specialty databases.

Protein families [40] were assigned, and then hypothetical proteins were identified. All proteins were then mapped to subsystems [41,42]. PubMLST (www.pubmlst.org) was used to assign sequence types, and then PhiSpy [43] was used to find prophages in bacterial genomes. Generated data includes genome quality assessment, antimicrobial resistance genes and phenotype predictions, specialty genes, subsystem overview, identification of the closest genome sequences, a phylogenetic tree, and a list of features that distinguish the genome from its most similar species.

### Comparing *Peptacetobacter hiranonis* Strains Cultured from the Feces of Dogs and Cats

The *P. hiranonis* genomes were compared using the average nucleotide identity (ANI), digital DNA-DNA hybridization (dDDH), and guanine-cytosine (GC) content. The ANI method with a pre-defined cutoff of 95% is widely accepted for species delineation, despite certain limitations [18–20]. In this study, we compared strains of *P. hiranonis* cultured from dogs and cats to determine whether those strains belong to (1) the same species, *P. hiranonis*, (2) different species, or even (3) different subspecies that may represent host-adapted bacterial subspecies.

The ANI was estimated by PYANI version 0.2.12 with “-m ANIm -g -v --workers 16” parameters, in which the percentage identity across all aligned regions, percentage coverage of each genome by aligned regions, number of bases from each genome contributing to the aligned regions, number of “similarity errors” on each genome, and a Hadamard matrix of percentage identity multiplied by percentage coverage for each comparison were generated [44]. A cutoff of 95% was defined for species delineation based on the percentage identity indicated by PYANI [18–20].

Digital DNA-DNA hybridization (dDDH) is an *in silico* method previously applied to the definition or delineation of species using wet-lab methodologies. The Type (Strain) Genome Server (TYGS: <u>Type Strain Genome Server</u>) applies the dDDH method to infer species and subspecies boundaries for prokaryotes, and it was employed for the genetic delineation of the *P. hiranonis* strains evaluated in this study [45,46]. For this, a predefined cutoff of 70% and 80% difference was used for species and subspecies delineation, respectively [21,22,46]. Additionally, differences in GC content on the *P. hiranonis* strains’ genomes were assessed – differences higher than 1% of GC content indicate different species.

### Phylogenetic Analysis of *Peptacetobacter hiranonis*

The Codon Tree pipeline generates a phylogenetic tree through the BV-BRC. In summary, to build the phylogenetic tree, the amino acid and nucleotide sequences from a defined number of global Protein Families were used [40], which were picked randomly to build an alignment, and then a tree based on the differences within those selected bacterial genomes, including the *P. hiranonis* strains. Protein sequences were aligned using MUSCLE [47], and the nucleotide coding gene sequences were aligned using the Codon Align function of BioPython [48]. The concatenated alignment of selected proteins and nucleotides was generated, describing the alignment [49]. Support values were generated using 1000 rounds of the “Rapid” bootstrapping option of RaxML [50].

### Biochemical Characterization of *Peptacetobacter hiranonis*

*P. hiranonis* strains (*n* = 22) were cultured in BBA plates (Anaerobe Systems, CA) in anaerobiosis at 37°C for 24 hours for the API^®^ tests (bioMérieux). A McFarland standard of 3.0 and 4.0 was used to prepare the inoculum for API^®^ 20A and API^®^ Rapid ID 32A, respectively, and tests were performed following the manufacturer’s instructions.

In summary, bacterial cells were harvested from BBA plates and suspended in the suspension medium (bioMérieux) as indicated by the manufacturer. The API^®^ strips, containing dehydrated substrates for the biochemical tests, were inoculated with the prepared bacterial suspension. The API^®^ 20A test was incubated in anaerobiosis at 37°C for 24 hours, while the API^®^ Rapid ID 32A test was incubated in aerobiosis at 37°C for 4 hours. After incubation, results were recorded as indicated by color changes in the API^®^ strip’s wells, and the characterization of *P. hiranonis* biochemical profile was recorded.

### Assessment of the *Peptacetobacter hiranonis*’ *in vitro* Ability to Deconjugate and Convert Bile Acids Using LC-MS/MS-Based Assay

The deconjugation of taurine-conjugated CA and CDCA (TCA and TCDCA) and conversion of primary (CA and CDCA) into secondary BA (DCA and LCA) and into oxo-BA were assessed using a liquid chromatography-tandem mass spectrometry (LC-MS/MS) based assay [51]. Strains were cultured for 48 hours in Brain-Heart Infusion (BHI) broth (Anaerobe Systems), and 250 µL of that culture was used as the initial inoculum for the deconjugation (*n =* 19) and conversion (*n =* 22) assays.

*P. hiranonis* were cultured in BHI broth supplemented with either 0.1 mM TCA, TCDCA (Cayman Chemical), CA, or CDCA (Thermo Scientific), and as a control in BHI without the addition of BA, for 24 hours at 37°C in anaerobiosis. After 24 hours of incubation of the BHI broth containing *P. hiranonis* and BA, the tube was thoroughly vortexed, and 100 µL of the suspension was transferred to a 2 mL screw-cap tube for BA extraction. Then, 300 µL of methanol was added to each tube, the tubes were shaken for 2 minutes using a Fisherbrand Bead Mill 24 (speed = 6 m/s, cycle = 1), and the tubes were centrifuged at 16,000 g for 10 minutes at 4°C.

Next, 200 µL of the supernatant was transferred to a microcentrifuge tube and centrifuged at 10,000 g for 5 minutes at 4°C. Finally, 150 µL of the resulting supernatant was transferred to a new 1.5 mL microcentrifuge tube and stored at –20°C until analysis. Prior to analysis, 20 µL internal standard mixture containing 0.02 mg/mL d_4_-GCA and 0.015 mg/mL d_4_-GLCA was added to each sample, vortexed, and 3 µL injected utilizing the analysis method previously described [51]. All strains were run in duplicate to assess the in vitro deconjugation of TCA and TCDCA, and in triplicate to assess the *in vitro* conversion of CA and CDCA. Culture without the addition of BA was performed as a blank sample for each strain tested, and the BA-converting *Clostridium scindens* (wild-type strain) was used as a control for assessing conversion.

Following extraction, conjugated, primary, secondary, oxo-, and iso-BA were then quantified using LC-MS/MS. This assay allows the quantification of the following BA: TCA, TCDCA, CA, CDCA, DCA, LCA, 12oxo-lithocholic (12oxo-LCA), 3oxo-cholic (3oxo-CA), 3oxo-chenodeoxycholic (3oxo-CDCA), 3oxo-deoxycholic (3oxo-DCA), 7,12oxo-lithocholic (7,12oxo-LCA), 7oxo-deoxycholic (7oxo-DCA), 7oxo-lithocholic (7oxo-LCA), glycocholic (GCA), glycochenodeoxycholic (GCDCA), glycodeoxycholic (GDCA), glycolithocholic (GLCA), glycoursodeoxycholic (GUDCA), hyocholic (HCA), hyodeoxycholic (HDCA), iso-cholic (Iso-CA), iso-chenodeoxycholic (Iso-CDCA), taurodeoxycholic acid (TDCA), taurolithocholic (TLCA), tauroursodeoxycholic (TUDCA), ursocholic (UCA), and ursodeoxycholic (UDCA) acids. Results were converted to micromolar concentrations and reported as % of total µM bile acids measured.

### Assessment of the Antimicrobial Susceptibility Profile of *Peptacetobacter hiranonis*

Although there is an agar dilution method for antimicrobial susceptibility testing for anaerobic bacteria, we opted to use the ETEST method in our study for practical reasons [52]. We recognized this as a limitation of our study, as ETEST^®^ is not recognized as a CLSI^®^ standard method. The ETEST^®^ method was used to estimate the minimum inhibitory concentration (MIC) for 11 *P. hiranonis* strains cultured from dogs and 11 strains from cats. Only one strain per animal was tested. In addition, the MIC concentration was also defined for the *P. hiranonis* DSM 13275, a reference strain.

For this, *P. hiranonis* strains were cultured in BBA (Anaerobe Systems) for 24 hours at 37°C in anaerobiosis. Colonies were harvested from BBA and transferred to *Brucella* broth (Anaerobe Systems, CA) at a concentration equivalent to McFarland 1.0. Cotton swabs were used to inoculate the bacterial suspension into BBA (Anaerobe Systems), and the plastic ETEST^®^ strips containing an antimicrobial in different concentrations were placed in the center of each plate. The inhibition zone and MIC were recorded after 24 hours of incubation.

In total, eighteen antimicrobials were tested: amoxicillin (antimicrobial concentration range: 0.016 – 256 µg/mL), amoxicillin/clavulanate (0.016 – 256), azithromycin (0.016 – 256), cefepime (0.016 – 256), cefoxitin (0.016 – 256), ceftriaxone (0.002 – 32), cephalothin (0.016 – 256), chloramphenicol (0.016 – 256), ciprofloxacin (0.002 – 32), clindamycin (0.016 – 256), daptomycin (0.016 – 256), meropenem (0.002 – 32), metronidazole (0.016 – 256), rifampicin (0.016 – 256), streptomycin (0.064 – 1024), tetracycline (0.016 – 256), trimethoprim/sulfamethoxazole (0.002 – 32), and vancomycin (0.016 – 256).

In addition, genes correlated to antibiotic resistance were assessed using the Comprehensive Genome Analysis through BV-BRC. In summary, genes with homology to those identified as being reported previously involved in antimicrobial resistance were blasted against proteins from the Comprehensive Antibiotic Resistance Database (CARD) [53], the National Database of Antibiotic-Resistant Organisms [54], the Antibiotic Resistance Database [55], and a curated database by BV-BRC [56]. Genes homologous to transporters were identified by searching against proteins from the Transporter Classification Database [57], and those similar to genes identified as potential drug targets by comparison with proteins from DrugBank [58] and the Therapeutic Target Database [59].

## RESULTS AND DISCUSSION

### Identification of *Peptacetobacter hiranonis* and Detection of BSH and *baiCD* Genes

In total, 11 *P. hiranonis* strains cultured from dog feces and 11 strains from cat feces were selected for further characterization. All *P. hiranonis* strains tested positive for *P. hiranonis’* 16S ribosomal RNA gene. The BSH gene, encoding the enzyme that deconjugates glycine- and taurine-conjugated BA, was detected in 90.9% (10/11) of the canine-derived *P. hiranonis*, except for the strain CH2. While the *baiCD* gene, one of the genes that belong to the bai operon responsible for the conversion of primary into secondary BA, was detected in 100% (11/11) of the canine-derived *P. hiranonis.* In the feline-derived strains, the BSH gene was detected in 18.2% (2/11) of the strains (CH5 and CH23), while the *baiCD* gene was detected in 90.9% (10/11) of the strains except for the strain CH18 (Table 2).

**Table 2.** Detection of the bile salt hydrolase (BSH) gene – gene encoding the enzyme that deconjugates glycine- and taurine-conjugated bile acids – and a bile acid-inducible (bai) operon gene, *baiCD* – one of the genes that belongs to the bai operon, responsible for the conversion of primary into secondary unconjugated bile acids – in feline- and canine-derived strains of *Peptacetobacter hiranonis*.

|  | **Bile Salt Hydrolase Gene** | **Bile Acid-Inducible CD (*baiCD*)** |
| --- | --- | --- |
|  | Deconjugation of Bile Acids | Conversion of Bile Acids |
| **Canine-derived *P. hiranonis*** |  |  |
| Detected | 10/11 (90.9%) | 11/11 (100%) |
| **Feline-derived *P. hiranonis*** |  |  |
| Detected | 2/11 (18.2%) | 10/11 (90.9%) |

### Whole-Genome Analysis of *Peptacetobacter hiranonis*

The comprehensive genome analysis of 21 strains of *P. hiranonis* is reported in Figure 1 and Supplementary Table 2. The genomes were assembled into a single contig; *P. hiranonis* strains have a total length ranging from 2,419,787 to 2,869,056 bp and G+C content ranging from 31.26 to 31.55%. The number of predicted protein-coding genes ranged from 2,128 to 2,552. The number of genes that were assigned to putative function ranged from 1,210 to 1,328.

**Figure 1.**
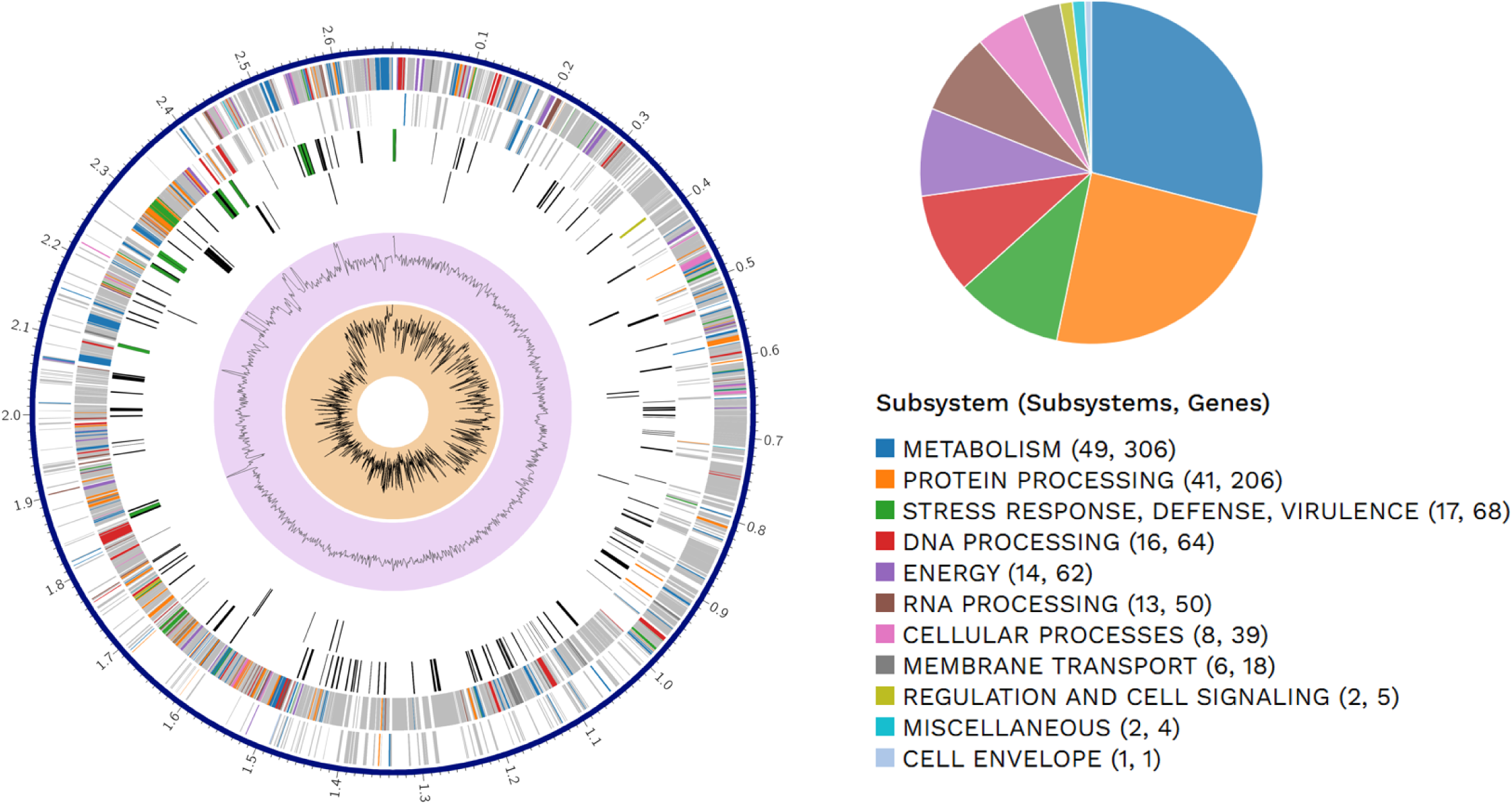
The circular genome of *Peptacetobacter hiranonis* strain CH2 is illustrated, providing the genome annotations. The circular genome includes, from outer to inner rings, contigs, forward and reverse strand coding sequences, RNA genes, coding sequences with homology to known antimicrobial genes, coding sequences with homology to known virulence factors, GC content, and GC skew. The colors of the coding sequences indicate the subsystem to which genes belong.

Coarse consistency, indicating the reliability of the annotation, ranged from 93.1 to 96.8%. Completeness of the evaluated genomes ranged from 76.6 to 82.6%; this measurement indicates the percentage of universal roles, which are usually found in a given taxonomic group, that are present in the evaluated genomes. Contamination ranged from 0 to 0.7%. The previously mentioned parameters, including contig count, DNA size, contigs N50 and L50, overrepresented, underrepresented, predicted, and total distinct roles, protein-encoding genes with and without functional assignment, protein-encoding feature coverage, hypothetical features, and genome length, can be found in Supplementary Table 3.

### Phylogenetic Analysis of *Peptacetobacter hiranonis*

Based on the clustering of genomes in the phylogenetic analysis of *P. hiranonis* isolated from fecal samples of dogs and cats, it is possible to observe that two separate clades are formed, one dominated by canine-derived strains and a second clade comprised mostly of feline-derived strains. These phylograms, shown in Figures 2 and 3, were built based on the 16S rRNA gene using RaxML and the whole genome using Genome BLAST Distance Phylogeny (GBDP) of the strains evaluated.

**Figure 2.**
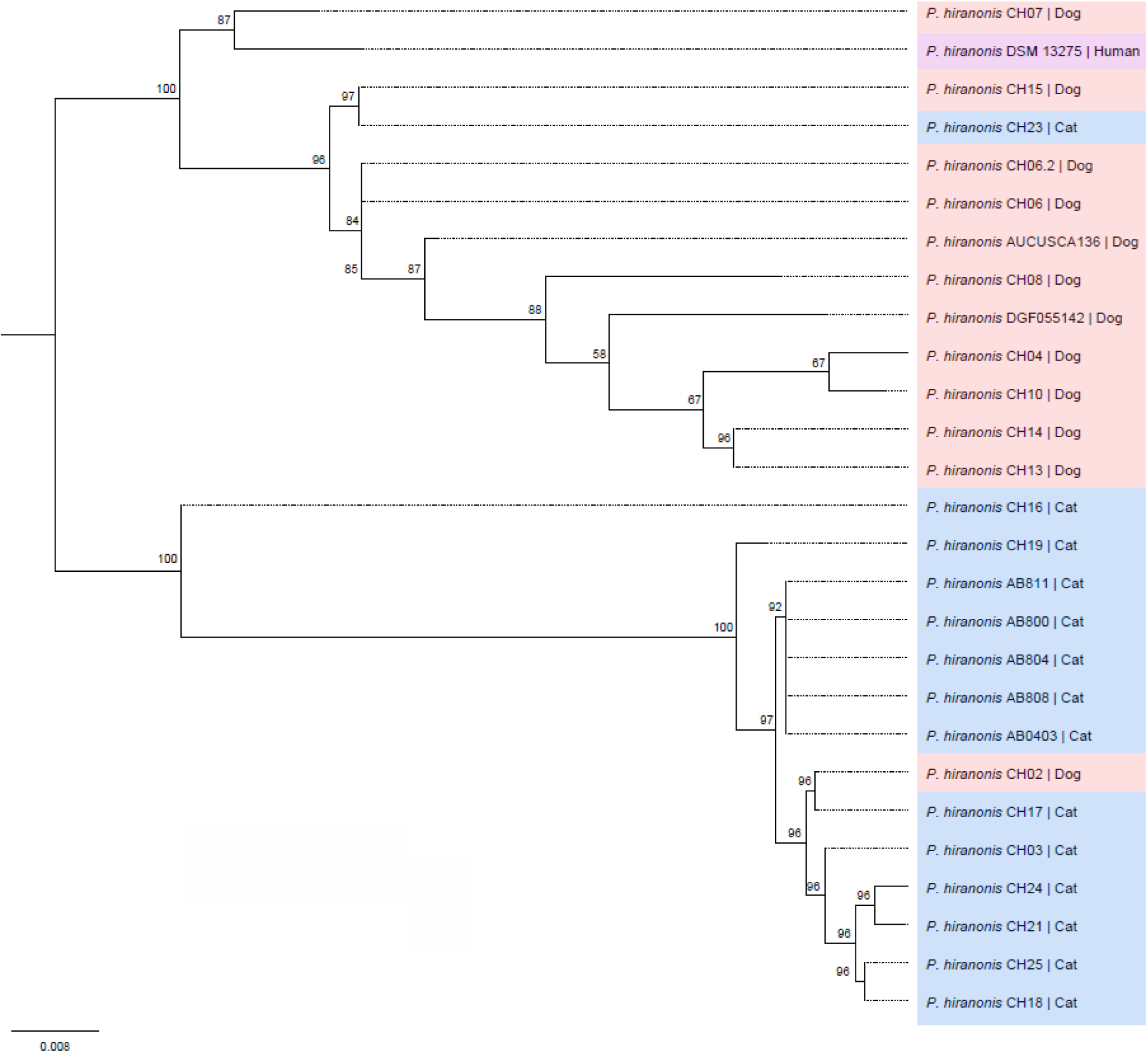
Maximum-likelihood tree showing the phylogenetic relationships of *Peptacetobacter hiranonis* strains described in this study and publicly available isolates from dogs (light red) and cats (light blue), and the reference strain *P. hiranonis* DSM 13275 (light purple), based on 16S rRNA gene sequences. Bootstrap percentages (≥50 %) based on 1000 resamplings are shown at the nodes. Abbreviations: CH#: *P. hiranonis* strains cultured from dogs and cats and described in the present study AB#, AUCUSCA136, and DGF055142: *P. hiranonis* strains cultured from cats and dogs whose genomes are publicly available.

**Figure 3.**
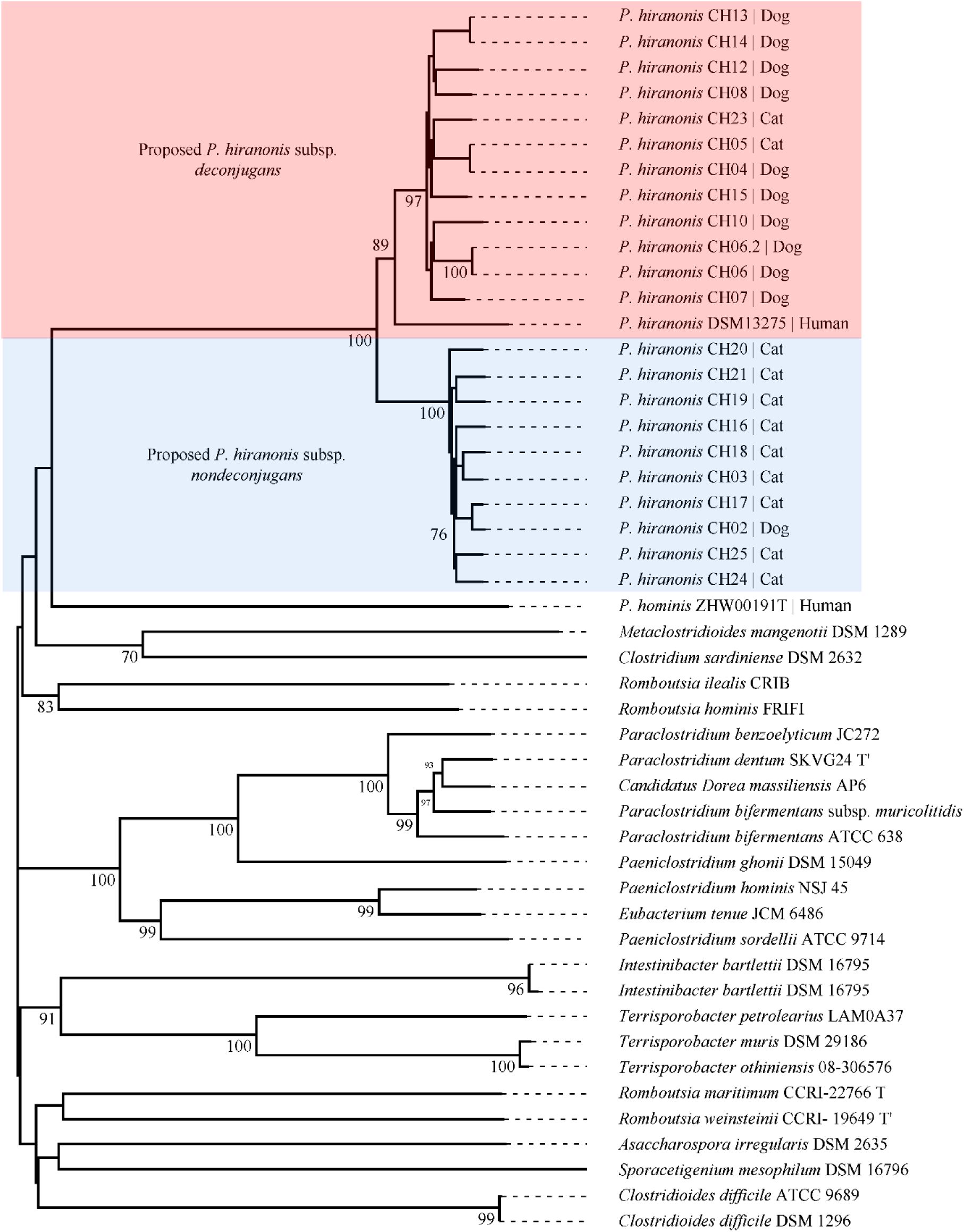
Whole-genome phylogenetic analysis of bile acid-converting bacterium *Peptacetobacter hiranonis* strains. The highlighted clades indicate the two proposed subspecies-level lineages within *P. hiranonis*: subsp. *deconjugans* in light red and subsp. *nondeconjugans* in light blue. *P. hiranonis* subsp. *deconjugans* includes the *P. hiranonis* DSM 13275 reference strain. Species and subspecies assignments were based on genome similarity analyses, including dDDH and ANI values, rather than on phylogenetic topology alone. The tree was inferred with FastME v2.1.6.1 using GBDP distances calculated from genome sequences. Branch lengths are scaled according to the GBDP distance formula d5. Numbers above branches indicate GBDP pseudo-bootstrap support values >60% based on 100 replications; the average branch support was 57.8%. The tree was rooted at the midpoint.

Interestingly, despite the BSH gene not being detected in 10/22 *P. hiranonis* strains and therefore not being included in the construction of the phylogenetic tree, strains were grouped based on their BA deconjugation ability. *P. hiranonis* CH2, the only canine-derived strain lacking the BSH gene, is grouped with the nine feline-derived strains that also lack the BSH gene. In addition, the two feline-derived strains, CH5 and CH23, which carry the BSH gene responsible for BA deconjugation, are grouped with the ten canine-derived *P. hiranonis* strains that also carry the BSH gene (Figures 2 and 3).

The association between phylogenetic lineages and BA deconjugation ability was observed in the strains evaluated. All strains in which the BSH gene was detected, or BA-deconjugating *P. hiranonis* strains, cluster together in Figure 2 (clade composed of *P. hiranonis* strains CH4, CH6, CH7, CH8, CH10, CH13, CH14, CH15, CH23, DSM 13275, AUCUSCA136, and DGF055142).

Conversely, all strains in which the BSH gene was not detected, or BA-nondeconjugating strains, cluster together (clade composed of *P. hiranonis* strains CH2, CH3, CH16, CH17, CH18, CH19, CH21, CH24, CH25, and AB#). The canine-derived *P. hiranonis* strain CH2 and feline-derived *P. hiranonis* strains CH5 and CH23 are important crossover examples, suggesting that host origin alone did not define the lineages observed here.

Based on the concordance between phylogenetic clustering and BA deconjugation phenotype, we propose that these two lineages represent two subspecies-level groups within the bile acid-converting bacterium *P. hiranonis*. The BSH-positive cluster, which includes strains capable of deconjugating BAs and contains the reference strain *P. hiranonis* DSM 13275, is proposed as *P. hiranonis* subsp. *deconjugans*. The BSH-negative cluster, composed of strains lacking the BSH gene, is proposed as *P. hiranonis* subsp. *nondeconjugans*. These designations are used hereafter to refer to the two genomic and functional lineages.

The dDDH analysis supports genetic divergence between *P. hiranonis* subsp. *deconjugans* and *P. hiranonis* subsp. *nondeconjugans*. Using the dDDH to compare the strains carrying the BSH gene and those who do not carry the BSH gene to each other and to the *P. hiranonis* genome previously published cultured from a dog’s feces (GCF_016694795.1) used as reference genome, the similarity parameters d0 (length of all High Scoring Pairs (HSPs) divided by total genome length), d4 (sum of all identities found in HSPs divided by overall HSP length), and d6 (sum of all identities found in HSPs divided by total genome length) are described in Table 3 and Supplementary Figure 1 (reference comparison) and Supplementary Table 4 (all other comparisons).

**Table 3.** Genome-to-Genome Distance Comparison Between Canine- and Feline-Derived *Peptacetobacter hiranonis* Strains and the Reference Genome. This table presents the results of digital DNA–DNA hybridization (dDDH) analyses between canine- and feline- derived *Peptacetobacter hiranonis* query strains and the reference genome: *P. hiranonis* (GCF_016694795.1). Three dDDH formulas (d0, d4, and d6) and their corresponding confidence intervals (C.I.) are shown, along with the difference in G+C content between genomes. The additional comparisons between all strains to each other are presented in Supplementary Table 4.

| Query Strain | Proposed Subspecies | dDDH (d0, in %) | C.I. (d0, in %) | dDDH (d4, in %) | C.I. (d4, in %) | dDDH (d6, in %) | C.I. (d6, in %) | G+C content difference (in %) |
| --- | --- | --- | --- | --- | --- | --- | --- | --- |
| CH14 | *deconjugans* | 89.2 | (85.9 - 91.9) | 83.8 | (81.0 - 86.3) | 91.1 | (88.4 - 93.1) | 0.13 |
| CH13 | *deconjugans* | 89.2 | (85.9 - 91.9) | 83.8 | (81.0 - 86.3) | 91.1 | (88.4 - 93.1) | 0.13 |
| CH08 | *deconjugans* | 89.6 | (86.3 - 92.2) | 83.7 | (80.9 - 86.1) | 91.4 | (88.8 - 93.4) | 0.13 |
| CH05 | *deconjugans* | 79.7 | (75.7 - 83.1) | 83.2 | (80.4 - 85.7) | 83.2 | (79.9 - 86.1) | 0.04 |
| CH04 | *deconjugans* | 79.5 | (75.5 - 83.0) | 83.1 | (80.3 - 85.6) | 83 | (79.7 - 85.9) | 0.04 |
| CH15 | *deconjugans* | 87.1 | (83.6 - 90.0) | 82.2 | (79.3 - 84.7) | 89.1 | (86.3 - 91.5) | 0.02 |
| CH23 | *deconjugans* | 88.6 | (85.1 - 91.3) | 81.8 | (78.9 - 84.3) | 90.2 | (87.5 - 92.4) | 0.02 |
| CH12 | *deconjugans* | 76.1 | (72.2 - 79.7) | 81.7 | (78.9 - 84.3) | 79.9 | (76.5 - 83.0) | 0.06 |
| CH07 | *deconjugans* | 78 | (74.1 - 81.6) | 81.3 | (78.4 - 83.9) | 81.5 | (78.1 - 84.4) | 0.06 |
| CH06 | *deconjugans* | 84.7 | (80.9 - 87.8) | 81 | (78.1 - 83.6) | 87 | (84.0 - 89.5) | 0.09 |
| CH06.2 | *deconjugans* | 84.3 | (80.5 - 87.5) | 81 | (78.1 - 83.6) | 86.7 | (83.6 - 89.3) | 0.09 |
| CH10 | *deconjugans* | 70.7 | (66.8 - 74.4) | 78.4 | (75.5 - 81.1) | 74.6 | (71.1 - 77.8) | 0.05 |
| CH25 | *nondeconjugans* | 70.7 | (66.7 - 74.3) | 65.3 | (62.4 - 68.2) | 71.9 | (68.5 - 75.2) | 0.15 |
| CH21 | *nondeconjugans* | 74.2 | (70.2 - 77.8) | 64.4 | (61.5 - 67.2) | 74.8 | (71.3 - 78.0) | 0.09 |
| CH24 | *nondeconjugans* | 74 | (70.0 - 77.6) | 64.2 | (61.3 - 67.1) | 74.6 | (71.1 - 77.8) | 0.2 |
| CH19 | *nondeconjugans* | 76.3 | (72.3 - 79.9) | 64.2 | (61.3 - 67.0) | 76.6 | (73.1 - 79.7) | 0.03 |
| CH20 | *nondeconjugans* | 72.7 | (68.7 - 76.3) | 64 | (61.0 - 66.8) | 73.4 | (69.9 - 76.6) | 0.02 |
| CH02 | *nondeconjugans* | 73.7 | (69.7 - 77.3) | 63.8 | (60.9 - 66.7) | 74.2 | (70.8 - 77.4) | 0.15 |
| CH18 | *nondeconjugans* | 72.8 | (68.9 - 76.5) | 63.7 | (60.8 - 66.5) | 73.5 | (70.0 - 76.7) | 0.12 |
| CH17 | *nondeconjugans* | 73.9 | (69.9 - 77.5) | 63.5 | (60.6 - 66.4) | 74.4 | (70.9 - 77.6) | 0.19 |
| CH03 | *nondeconjugans* | 70.4 | (66.5 - 74.1) | 63.5 | (60.6 - 66.3) | 71.3 | (67.9 - 74.6) | 0.1 |
| CH16 | *nondeconjugans* | 77.8 | (73.8 - 81.3) | 63.4 | (60.5 - 66.3) | 77.7 | (74.2 - 80.8) | 0.12 |

We hence proposed that the two phylogenetic clades be reclassified as two distinct subspecies, namely *P. hiranonis* subsp. *nondeconjugans* and *P. hiranonis* subsp. *deconjugans*, based on the presence and absence of BA deconjugation abilities encoded by the BSH gene. The similarity scores for *P. hiranonis* subsp. *nondeconjugans* [median (range) d0: 74.3% (71.3 – 77.7); d4: 63.9% (63.4 – 65.3); and d6: 74.3% (71.3 – 77.7)] in comparison to the *P. hiranonis* reference genome were lower compared to the similarity scores of *P. hiranonis* subsp. *deconjugans* [d0: 86.8% (74.6 – 91.4); d4: 82% (78.4 – 83.8); and d6: 86.8% (74.6 – 91.4)] in comparison to the *P. hiranonis* reference genome.

The lower similarity scores obtained from the comparison between *P. hiranonis* subsp. *nondeconjugans* strains and the *P. hiranonis* reference genome (GCF_016694795.1) indicate greater divergence in the aligned genetic regions used for the TYGS analysis. Despite showing d4 similarity scores below 70%, d0 and d6 similarity scores are above 70%, which supports that they may not represent a new species but rather a subspecies as proposed above.

Through the analysis performed by PYANI evaluating the ANI percentage matrix – pairwise identity of all aligned regions – among all canine- and feline-derived strains of *P. hiranonis*, it was possible to observe that the ANI was higher than 95% across all *P. hiranonis* strains (Figure 4). Considering the alignment coverage matrix – reflecting the percentage of each genome covered by aligned regions – and the Hadamard matrix – the percentage identity multiplied by the percentage coverage for each comparison – percentages equal to or higher than 68% and 65% were found among all strains of *P. hiranonis*, respectively (Supplementary Figure 2).

**Figure 4.**
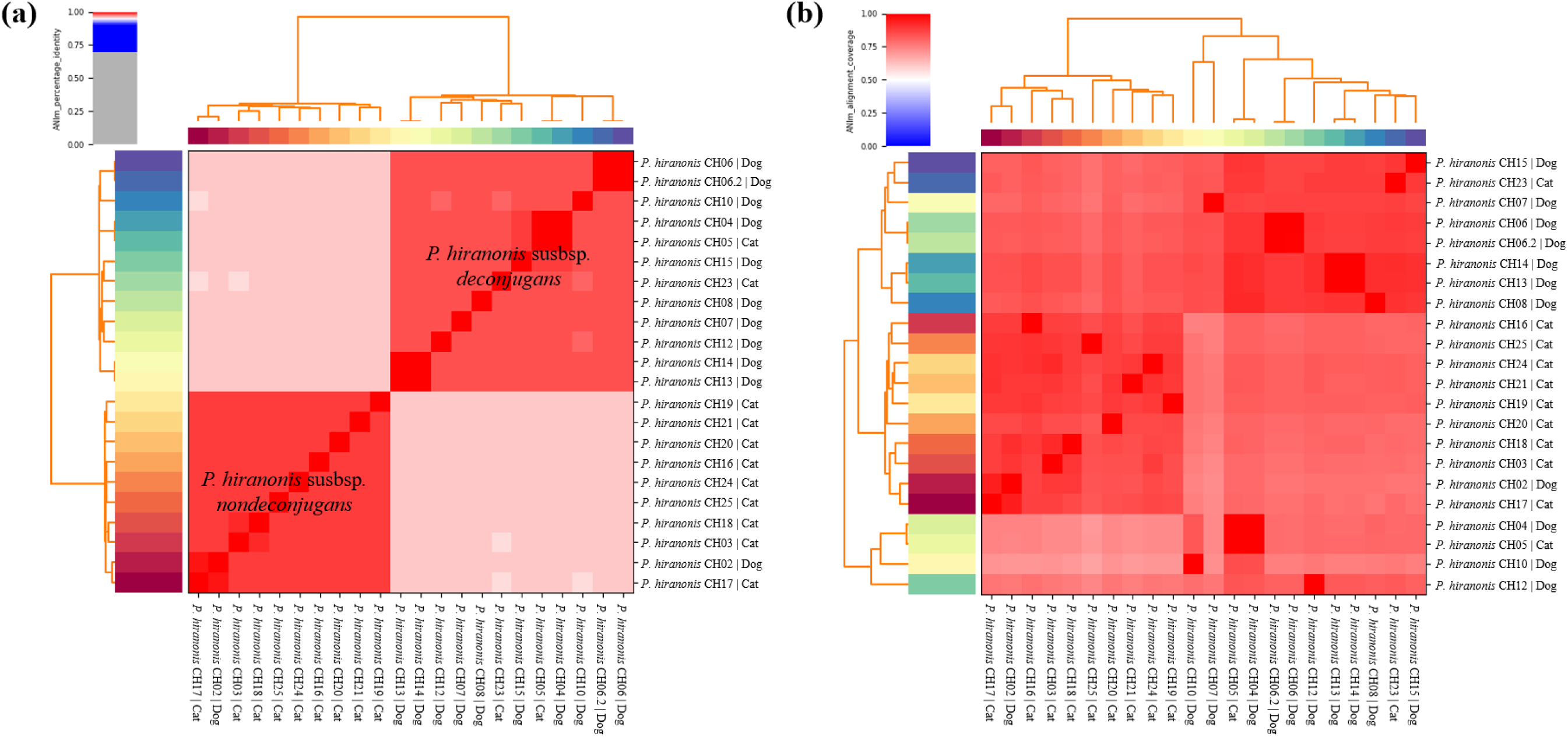
Percentage identity (a) and coverage (b) matrix for *Peptacetobacter hiranonis* ANIm analysis. Among all canine- and feline-derived strains of *P. hiranonis*, the percentage identity was greater than 95%. When comparing *P. hiranonis* subsp. *deconjugans* to subsp. *nondeconjugans*, the percentage identity was slightly lower, around 96%, but still above the 95% cutoff. Within each subspecies, the percentage identity ranged from 98% to 99%.

Based on the ANI results, two clusters were formed: one containing *P. hiranonis* strains carrying the BSH gene (*P. hiranonis* subsp. *deconjugans*), which included mostly strains cultured from dog feces, and a second cluster containing strains that did not carry the BSH gene (*P. hiranonis* subsp. *nondeconjugans*), mostly cultured from cat feces. When ANI is evaluated within clusters, the percentage identity ranges from 98% to 99%; in contrast, the ANI between clusters is 96% (Table 4).

**Table 4.** Proposed subspecies for the bile acid-converting bacterium *Peptacetobacter hiranonis:* subsp. *deconjugans* and subsp. *nondeconjugans*.

| Deconjugation Activity | Proposed Subspecies | Strain ID |
| --- | --- | --- |
| Absent | *P. hiranonis* subsp. *nondeconjugans* | CH2, CH3, CH16, CH17, CH18, CH19, CH20, CH21, CH24, and CH25 |
| Present | *P. hiranonis* subsp. *deconjugans* | CH4, CH5, CH6, CH7, CH8, CH10, CH12, CH13, CH14, CH15, and CH23 |

The dDDH and ANI analysis support genetic divergence between the two proposed subspecies, *P. hiranonis* subsp. *deconjugans* and *P. hiranonis* subsp. *nondeconjugans*; however, supporting keeping them within the same species, *P. hiranonis*. When comparing *P. hiranonis* subsp. *nondeconjugans* strains to the reference genome *P. hiranonis* (GCF_016694795.1) or to our proposed *P. hiranonis* subsp. *deconjugans* strains, the dDDH d4 value (see Table 4) is lower than the current proposed 70% cutoff for species delineation. In contrast, the dDDH d0 and dDDH d6 values are >70%, the ANI is > 95% (the proposed cutoff for species delineation using ANI), and the G+C content differences are < 1% (see Table 3; the proposed cutoff for species delineation\ using G+C content differences), strengthening the hypothesis that these strains do belong to the same species: *P. hiranonis*.

Interestingly, although the majority of *P. hiranonis* subsp. *deconjugans* strains were cultured from feces obtained from dogs, and the majority of *P. hiranonis* subsp.*nondeconjugans* strains were cultured from feces obtained from cats, a crossover between hosts was observed. Therefore, while a strong host preference was observed, we were unable to confirm host specificity for either *P. hiranonis* subsp. *deconjugans* and *P. hiranonis* subsp. *nondeconjugans* strains.

### Biochemical Profile of Peptacetobacter hiranonis subsp. deconjugans and subsp. nondeconjugans

*P. hiranonis* subsp. *deconjugans* (*n* = 12) and subsp. *nondeconjugans* (*n* = 10) strains were selected for biochemical profiling. Using API^®^ 20A to characterize *P. hiranonis* strains, positive reactions were observed on the acidification of broth by the utilization of glucose, saccharose, and mannose. Negative reactions were observed on the utilization of lactose, maltose, salicin, xylose, arabinose, gelatinase, glycerol, cellobiose, esculin, melibiose, raffinose, rhamnose, trehalose; and production of indole, urease, catalase, and hydrogen sulfide (H_2_S).

Variable results for the utilization of mannitol and sorbitol were observed. Five out of 10 strains of *P. hiranonis* subsp. *nondeconjugans* utilized sorbitol, highlighting that *P. hiranonis* strains in which utilization of sorbitol is detected may be classified presumptively as *P. hiranonis* subsp. *nondeconjugans* (Table 5 and Supplementary Table 5).

**Table 5.** Biochemical profiling of *Peptacetobacter hiranonis* subsp. *deconjugans* (n=12) and subsp. *nondeconjugans* (n=10) strains using API® 20A test.

| Biochemical Tests | Pos % | Neg % |
| --- | --- | --- |
| **Acidification from:** |  |  |
| Glucose, Saccharose and Mannose | 100 (22/22) | 0 (0/22) |
| Mannitol | 86.4 (19/22) | 13.6 (3/22) |
| Sorbitol | 22.7 (5/22) | 77.3 (17/22) |
| Indole formation, urease activity, acidification of lactose, maltose, salicin, xylose, arabinose, glycerol, melezitose, raffinose, rhamnose, trehalose, hydrolysis of gelatin and esculin, catalase, and production of H<sub>2</sub>S | 0 (0/22) | 100 (22/22) |

Abbreviations: Pos: positive reaction. Neg: negative reaction.

Using the API^®^ Rapid ID 32A test to characterize *P. hiranonis* subsp. *deconjugans* (*n*=12) and subsp. *nondeconjugans* (*n*=10) strains, positive reactions were observed in the production of N-Acetyl-ßglucosaminidase for all strains. Negative reactions were observed in the production of urease, α-galactosidase, ß-galactosidase, ß-galactosidase 6-phosphatase, α-glucosidase, ß-glucosidase, α-arabinosidase, ß-glucuronidase, mannose, raffinose, α-fucosidase, phenylalanine arylamidase, tyrosine arylamidase, glycine arylamidase, histidine arylamidase, glutamyl-glutamic acid arylamidase, and serine arylamidase.

Variable results in the production of glutamic acid decarboxylase, leucine arylamidase, pyroglutamic acid arylamidase, arginine arylamidase, alanine arylamidase, alkaline phosphatase, leucyl-glycine arylamidase, arginine dihydrolase, indole, and proline arylamidase, and reduction of nitrates were observed. Only feline-derived strains of *P. hiranonis* presented alanine arylamidase activity (Table 6 and Supplementary Table 5).

**Table 6.**
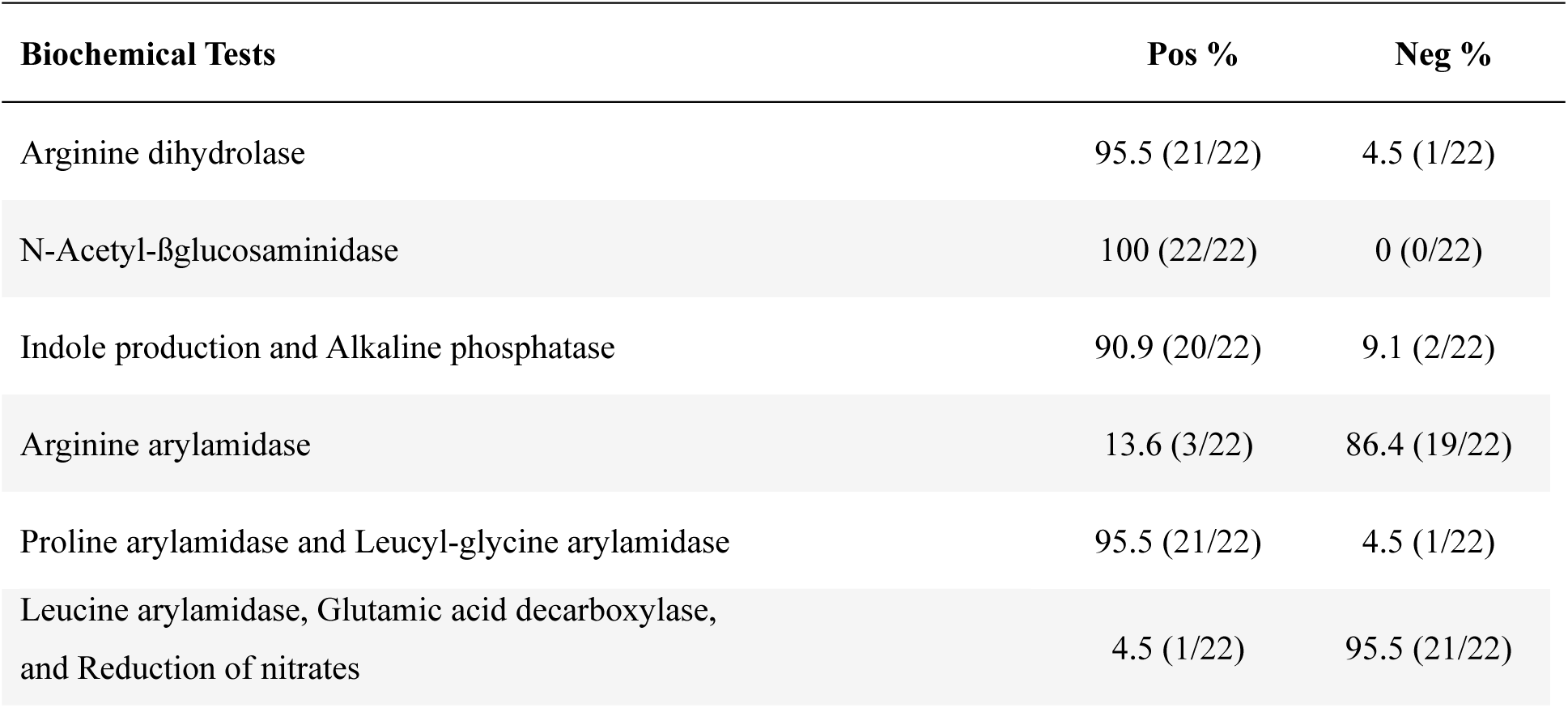
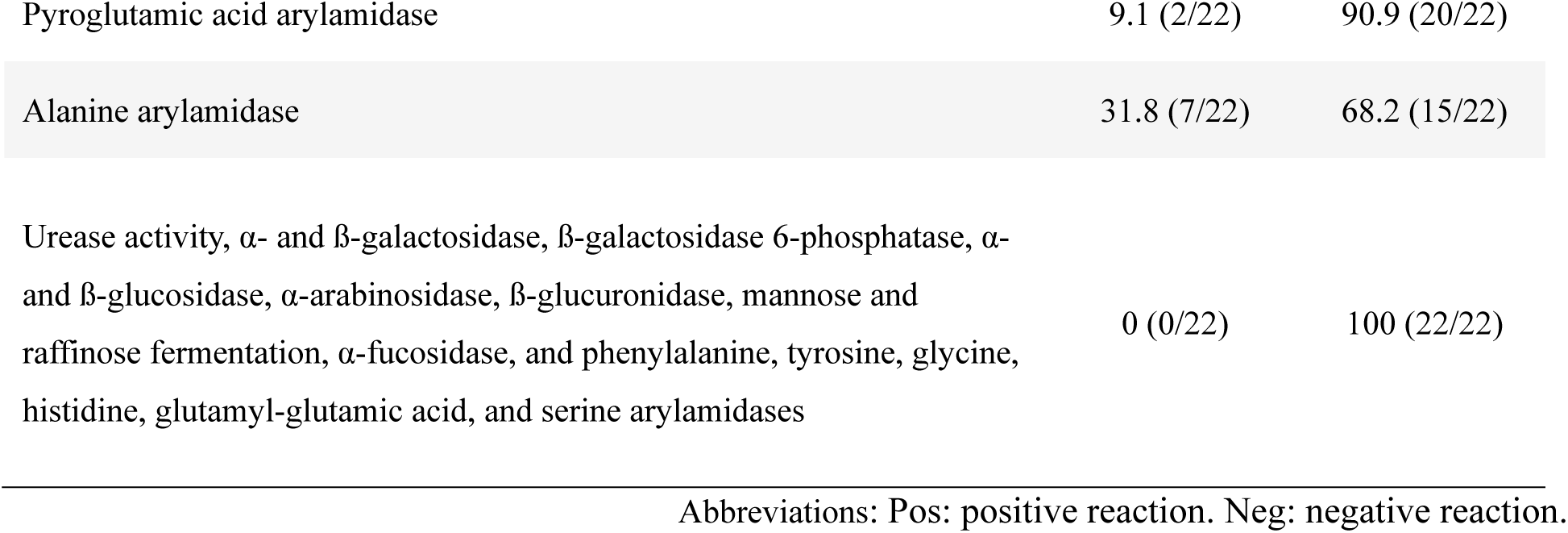
Biochemical profiling of *Peptacetobacter hiranonis* subsp. *deconjugans* (n = 12) and subsp. *nondeconjugans* (n = 10) using API® Rapid ID 32A test.

### Ability of *Peptacetobacter hiranonis* subsp. *deconjugans* and subsp. *nondeconjugans* to Deconjugate and Convert Bile Acids *In Vitro*

The deconjugation *in vitro* assay to evaluate the ability of strains to deconjugate taurine-conjugated cholic acid and chenodeoxycholic acid was performed in *P. hiranonis* subsp. *deconjugans* (*n=*10) and *P. hiranonis* subsp. *nondeconjugans* (*n=*9) strains. After 24 hours of incubation with TCA-spiked broth, TCA as a percentage of the total BAs measured ranged from 0 to 3% in the *P. hiranonis* subsp. *deconjugans* and from 95 to 96% in the *P. hiranonis* subsp. *nondeconjugans*.

After 24 hours of incubation with TCDCA-spiked broth, TCDCA as a percentage of the total BAs measured ranged from 0 to 12% in *P. hiranonis* subsp. *deconjugans* and from 97 to 99% in the *P. hiranonis* subsp. *nondeconjugans*. In summary, all *P. hiranonis* subsp. *deconjugans* strains were able to deconjugate TCA and TCDCA *in vitro* (Figure 5), and *P. hiranonis* subsp. *nondeconjugans* strains were unable to deconjugate TCA and TCDCA.

**Figure 5.**
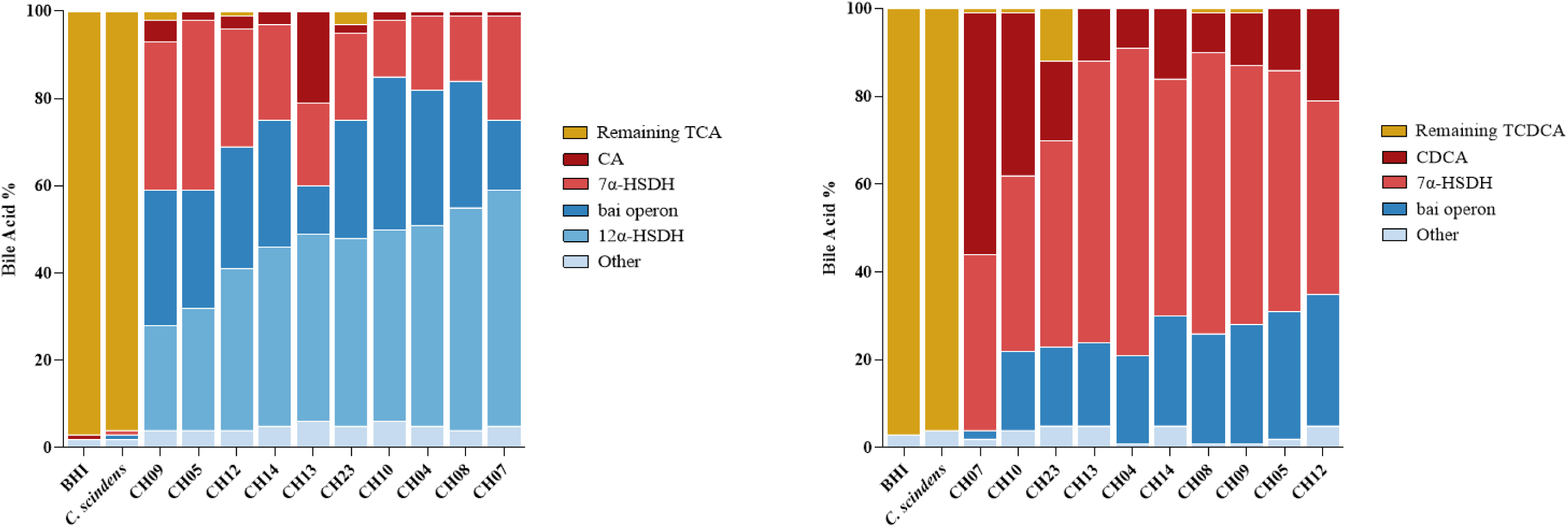
The in vitro ability of *Peptacetobacter hiranonis* subsp. *deconjugans* strains to deconjugate taurine-conjugated cholic acid (TCA, shown in A) and taurine-conjugated chenodeoxycholic acid (TCDCA, shown in B). *P. hiranonis* subsp. *nondeconjugans* strains were unable to deconjugate either TCA or TCDCA *in vitro*; hence, they are not represented here. *Clostridium scindens* was used as a negative control, as it cannot deconjugate bile acids. Negative controls (Brain-Heart Infusion (BHI) broth spiked with TCA or TCDCA but not inoculated with bacteria) are represented in the graph. Abbreviations: TCA: Taurine-Conjugated Cholic Acid. TCDCA: Taurine-Conjugated Chenodeoxycholic Acid. CA: Cholic Acid. CDCA: Chenodeoxycholic Acid. bai: Bile Acid-Inducible. HSDH: Hydroxysteroid Dehydrogenase.

The bai operon activity was noted by the presence of 7α-dehydroxylation products of CA and CDCA: DCA and LCA, respectively. Oxidation of the primary and secondary BAs’ hydroxyl groups by a position-specific hydroxysteroid dehydrogenase (HSDH) also occurred when evaluating the *in vitro* deconjugation and conversion ability of *P. hiranonis* for both CA and CDCA. For CA, *P. hiranonis* strains were capable of oxidizing CA into 7-oxo-DCA, and DCA into 12-oxo-LCA, using 7α-HSDH and 12α-HSDH, respectively. For CDCA, *P. hiranonis* strains were capable of oxidizing CDCA into 7-oxo-LCA using a 7α-HSDH (Figures 5, 6, and 7).

We next evaluated 7α-dehydroxylation ability (also referred to as conversion of primary into secondary unconjugated BAs) by spiking with unconjugated primary BAs, CA, or CDCA. In total, 12 strains of *P. hiranonis* subsp. *deconjugans* and 10 strains of *P. hiranonis* subsp. *nondeconjugans* were evaluated. The remaining spiked CA, which was not converted by *P. hiranonis,* ranged from 1 to 59% (median: 9.5%) of the total BAs measured.

*P. hiranonis* strains converted CA into 7-oxo-DCA, median (range): 18.5% (1 to 29%) of total BA measured, through the 7α-hydroxysteroid dehydrogenase enzyme (HSDH); and into DCA and HSDH products of DCA, median (range): 70% (0 to 92%) of total BA measured, through the BAI operon. In addition, conversion of DCA into 12-oxo-LCA, median (range): 28% (0 to 63%) of total BA measured through the 12α-HSDH was also observed *in vitro* (Figure 6).

**Figure 6.**
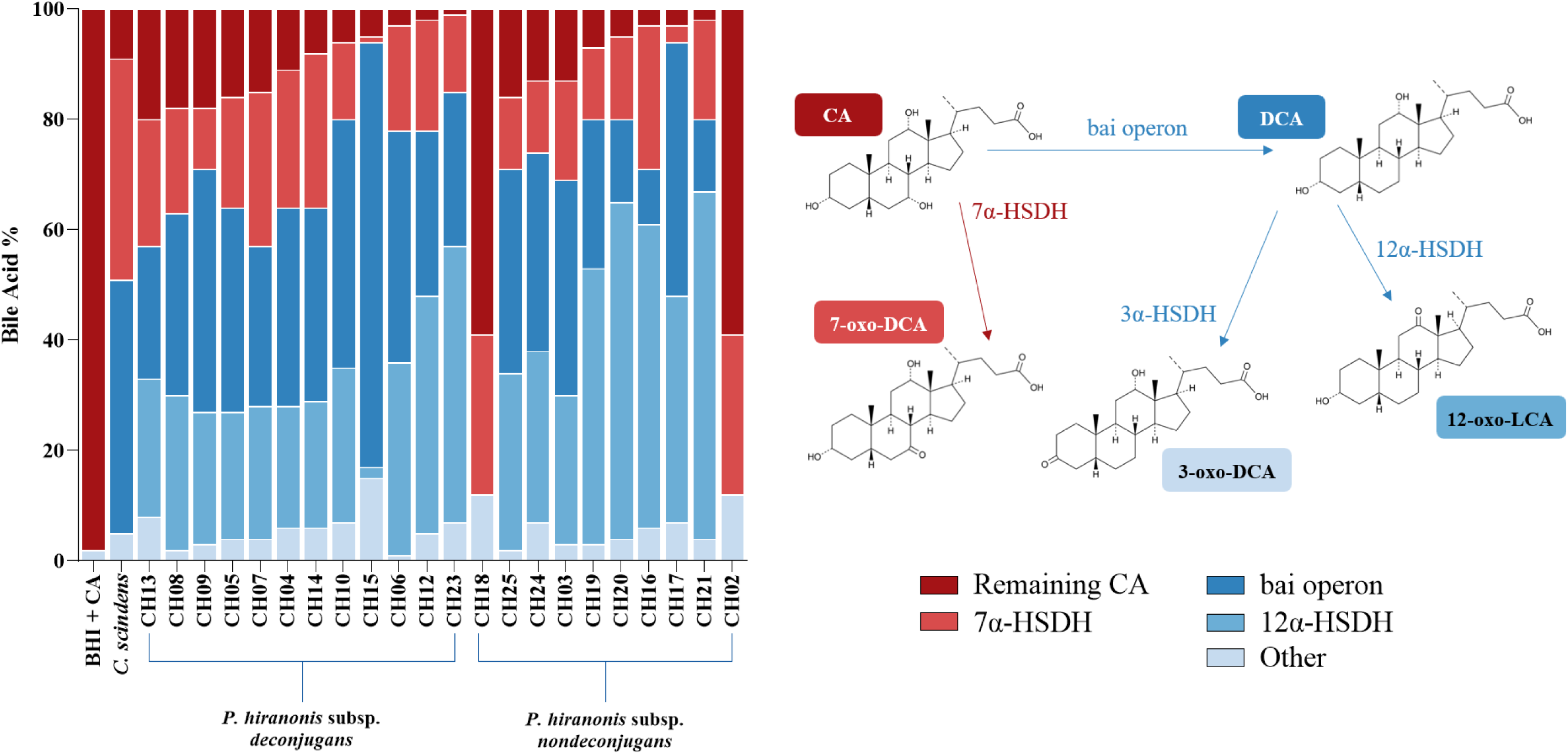
*Peptacetobacter hiranonis* subsp. *deconjugans* and subsp. *nondeconjugans* strains’ ability to convert CA into DCA using the BAI operon; to convert CA into 7-oxo-DCA using a 7α-HSDH; to convert DCA into 3-oxo-DCA using a 3α-HSDH; and to convert DCA into 12-oxo-LCA using a 12α-HSDH. CA: Cholic Acid. DCA: Deoxycholic Acid. BAI: Bile Acid-Inducible. HSDH: Hydroxysteroid Dehydrogenase. LCA: Lithocholic Acid. BHI: Brain-Heart Infusion Broth.

The remaining spiked CDCA, which was not converted by *P. hiranonis,* ranged from 3 to 71% (median: 27%) of the total BA measured. Strains converted CDCA into 7-oxo-LCA, median (range): 56% (5 to 84) of total BA measured, through the 7α-HSDH; and into LCA, median (range): 15% (0 to 30) of total BA measured, through the BAI operon (Figure 7). No differences were observed in the ability of *Peptacetobacter hiranonis* subsp. *deconjugans* and subsp. *nondeconjugans* strains when comparing the conversion of both CA and CDCA into DCA and LCA, respectively. However, the two strains that did not 7α-dehydroxylate CA, canine CH2 and feline CH18, were the same strains that did not 7α-dehydroxylate CDCA, and both of these strains were *Peptacetobacter hiranonis* subsp. *nondeconjugans*. Interestingly, the presence or absence of the *baiCD* gene via qPCR matched the strains’ *in vitro* 7α-dehydroxylation activity in all strains except canine CH2, where the gene was present, but the activity was not.

**Figure 7.**
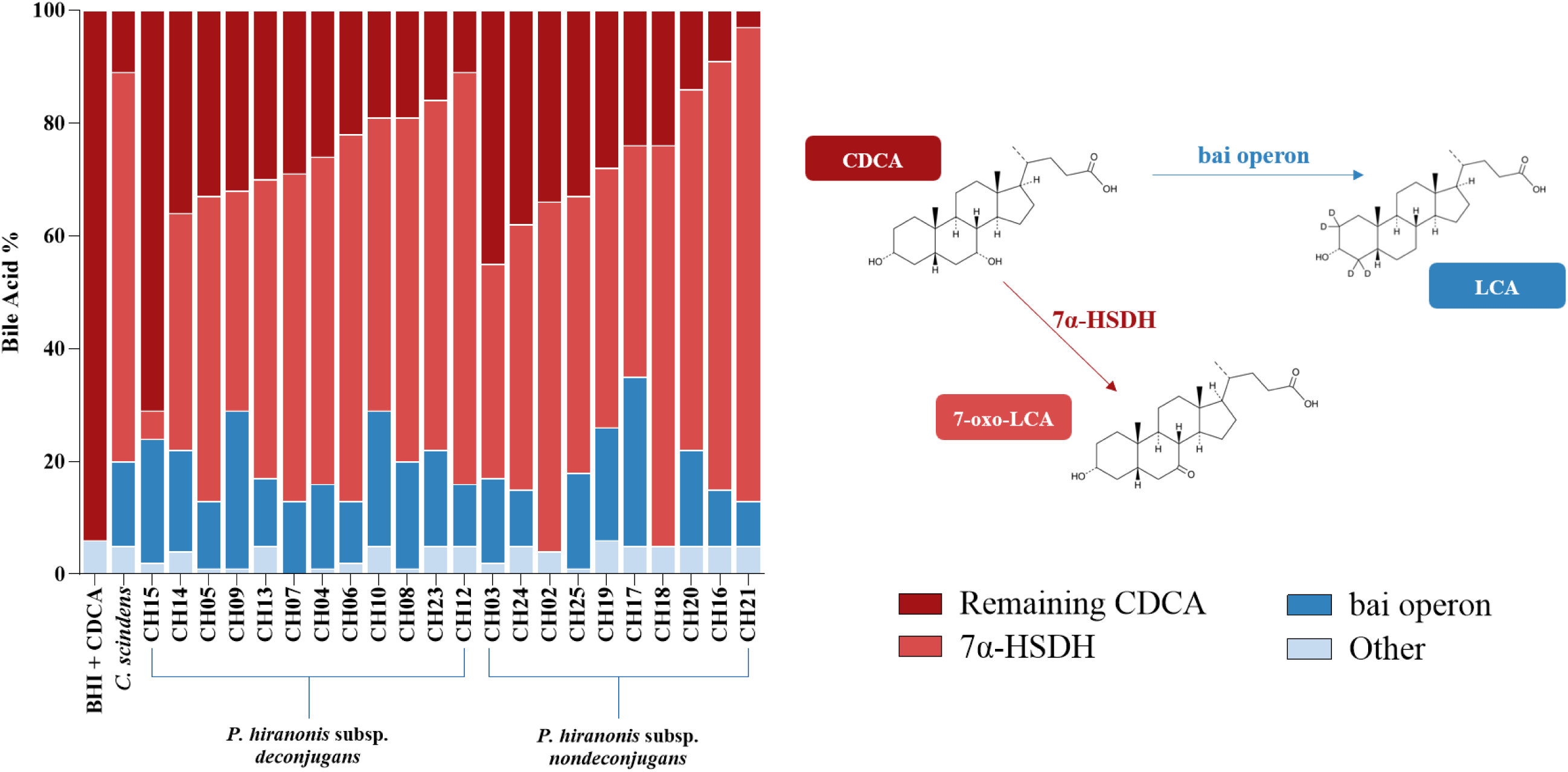
*Peptacetobacter hiranonis* subsp. *deconjugans* and subsp. *nondeconjugans* strains’ ability to convert CDCA into LCA using the BAI operon, and to convert CDCA into 7-oxo-LCA using a 7α-HSDH. CDCA: Chenodeoxycholic Acid. LCA: Lithocholic Acid. BAI: Bile Acid-Inducible. HSDH: Hydroxysteroid Dehydrogenase. BHI: Brain-Heart Infusion Broth

After the 24-hour incubation period, more DCA was produced from CA relative to LCA production from CDCA. This may suggest enzymatic preference for bile acids containing a 12-hydroxyl group. Shorter or longer incubation periods were not tested but may help reveal kinetics of the bile acid deconjugations and conversions.

Individual variability in the ability to deconjugate TCA and TCDCA, as well as to convert primary into secondary unconjugated BAs, was observed among *P. hiranonis* subsp. *deconjugans* and subsp. *nondeconjugans* strains. However, currently, we do not have evidence that differences in the ability to deconjugate or convert BAs influence canine or feline intestinal health, nor that these differences in the ability to deconjugate or convert BAs among strains have a significant effect on the BA acid metabolism of companion animals presenting with microbial BA dysmetabolism.

It is important to highlight that the presence or absence of deconjugation and conversion abilities, although variable in their intensity, is likely more clinically relevant than focusing solely on differences in deconjugation and conversion percentages across *P. hiranonis* strains. Moreover, as with other bacteria, such as *Staphylococcus* found on skin or *Escherichia coli* found in the intestines, animals may simultaneously harbor multiple strains of *P. hiranonis*. In this study, however, we described only one strain cultured from each dog or cat.

Additionally, it is important to note the limitations of using an *in vitro* model, which inherently cannot reflect all the variables present in an *in vivo* environment. *In vivo, P. hiranonis* must compete for nutrients with other bacteria present in the intestinal microbiota and is constantly exposed to pressure from the host immune system; factors that may, in part, influence the ability of these strains to deconjugate and convert BAs.

Summary table presenting mean values for bile acid deconjugation and conversion, as well as the presence or absence of *bai* operon genes in *Peptacetobacter hiranonis* subsp. *deconjugans* and subsp. *nondeconjugans* can be found in Supplementary Table 7.

### Antimicrobial Susceptibility Profiling of *Peptacetobacter hiranonis* subsp. *deconjugans* and subsp. *nondeconjugans* Strains

An MIC of ≤3 µg/mL for amoxicillin, amoxicillin/clavulanate, cefepime, ceftriaxone, chloramphenicol, ciprofloxacin, clindamycin, daptomycin, meropenem, metronidazole, rifampicin, and vancomycin inhibited the growth of all *Peptacetobacter hiranonis* subsp. *deconjugans* and subsp. *nondeconjugans* strains. Trimethoprim and sulfamethoxazole did not inhibit the growth of any tested strains at a concentration of 32 µg/mL (see Table 7, Figure 8, and Supplementary Table 6). No differences were observed in the antimicrobial susceptibility profile when comparing either canine- to feline-derived *P. hiranonis* strains or *Peptacetobacter hiranonis* subsp. *deconjugans* to subsp. *nondeconjugans*.

**Figure 8.**
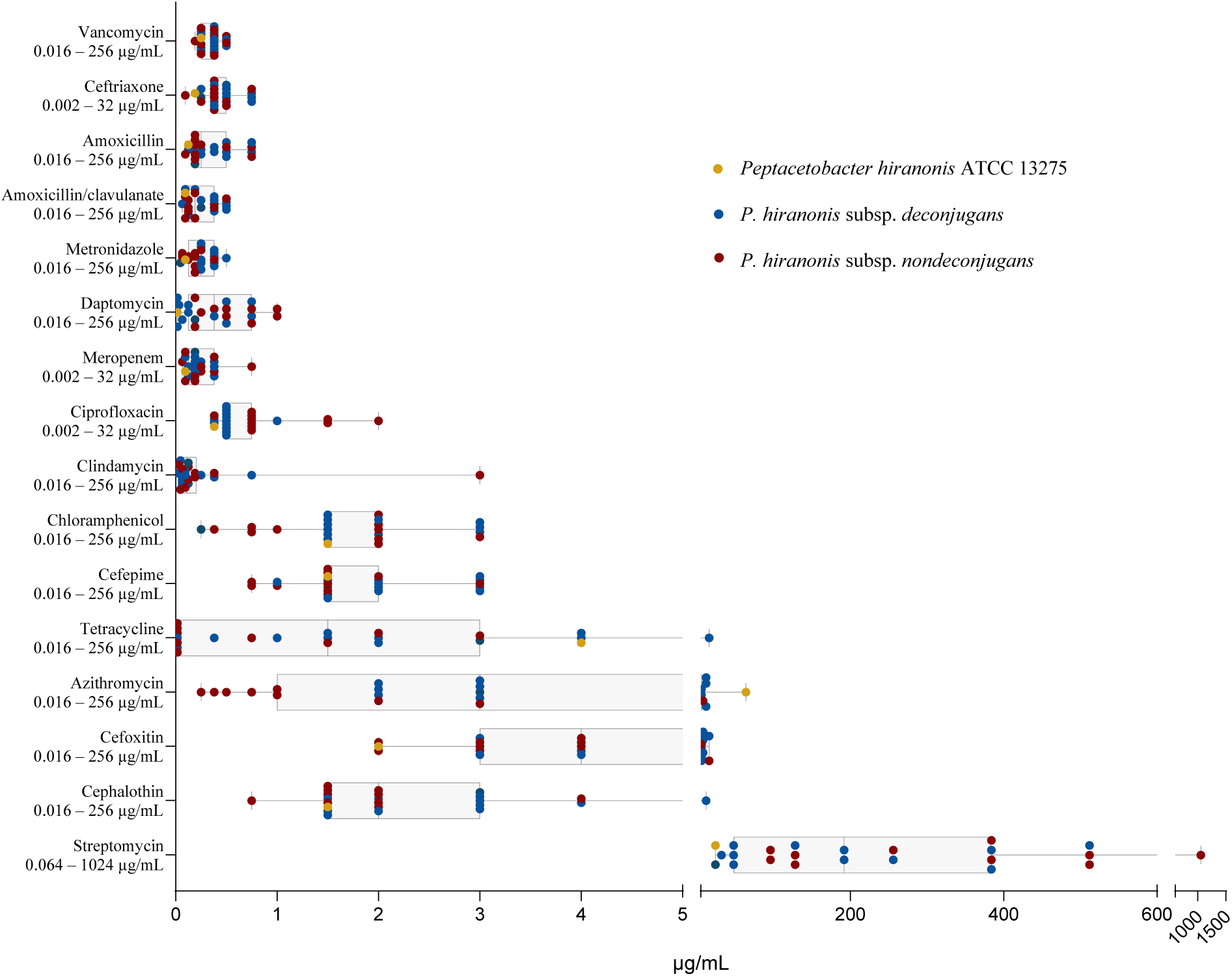
Minimum Inhibitory Concentration for *Peptacetobacter hiranonis* subsp. *deconjugans* and subsp. *nondeconjugans* strains assessed using ETEST®. Note: Trimethoprim and sulfamethoxazole did not inhibit the growth of any tested strains at a maximum concentration of 32 µg/mL, and it is not displayed in this figure.

**Table 7.** Median, minimum, and maximum values for the minimum inhibitory concentration assessed using ETEST® in strains of *Peptacetobacter hiranonis*. No significant differences were observed between canine- and feline-derived or *Peptacetobacter hiranonis* subsp. *deconjugans* to subsp. *nondeconjugans* strains.

| Antimicrobial<br>Concentration Range | ETEST Results *n* = 22<br>Median (Min – Max) | *P. hiranonis*<br>DSM 13275 |
| --- | --- | --- |
| Amoxicillin 0.016 – 256 | 0.25 (0.094 – 0.75) | 0.125 |
| Amoxicillin/clavulanate 0.016 – 256 | 0.22 (0.064 – 0.5) | 0.094 |
| Azithromycin 0.016 – 256 | 3 (0.25 – >256) | 64 |
| Cefepime 0.002 – 256 | 1.5 (0.75 – 3) | 1.5 |
| Cefoxitin 0.016 – 256 | 4 (2 – 16) | 2 |
| Ceftriaxone 0.002 – 32 | 0.38 (0.094 – 0.75) | 0.19 |
| Cephalothin 0.016 – 256 | 2 (0.75 – 12) | 1.5 |
| Chloramphenicol 0.016 – 256 | 1.75 (0.25 – 3) | 1.5 |
| Ciprofloxacin 0.002 – 32 | 0.5 (0.38 – 2) | 0.380 |
| Clindamycin 0.016 – 256 | 0.1095 (0.023 – 3) | >256 |
| Daptomycin 0.016 – 256 | 0.38 (0.016 – 1) | 0.016 |
| Meropenem 0.002 – 32 | 0.19 (0.064 – 0.75) | 0.094 |
| Metronidazole 0.016 – 256 | 0.25 (0.047 – 0.5) | 0.094 |
| Rifampicin 0.016 – 256 | 0.002 (0.002 – 0.5) | 0.002 |
| Streptomycin 0.064 – 1024 | 192 (24 – >1024) | 24 |
| Tetracycline 0.016 – 256 | 1.5 (0.016 – >256) | 4 |
| Trimethoprim/Sulfamethoxazole 0.002 – 32 | Not inhibited (>32) | >32 |
| Vancomycin 0.016 – 256 | 0.38 (0.19 – 0.5) | 0.25 |

#### Antimicrobial Resistance Genes

The antimicrobial resistance genes (AMR) are presented in Table 8, and they are divided as (1) antibiotic target in susceptible species, (2) antibiotic target-modifying enzyme, (3) antibiotic target replacement protein, (4) efflux pump conferring antibiotic resistance, (5) gene conferring resistance via the absence, or (6) protein-altering cell wall charge conferring antibiotic resistance. Although AMR genes were detected, the *in vitro* susceptibility assessed using E-TEST^®^ indicates that tested strains were inhibited at low concentrations of amoxicillin [median (min – max) MIC: 0.25 (0.094 – 0.75)], amoxicillin/clavulanate [(0.22 (0.064 – 0.5)], cefepime [1.5 (0.75 – 3)], ceftriaxone [0.38 (0.094 – 0.75)], chloramphenicol [1.75 (0.25 – 3)], ciprofloxacin [0.5 (0.38 – 2)], clindamycin [0.1095 (0.023 – 3)], daptomycin [0.38 (0.016 – 1)], meropenem [0.19 (0.064 – 0.75)], metronidazole [0.25 (0.047 – 0.5)], rifampicin [0.002 (0.002 – 0.5)], and vancomycin [0.38 (0.19 – 0.5)]. Inhibition by trimethoprim-sulfamethoxazole at a concentration equal to 32 µg/ml was not observed (see Table 7, and Supplementary Figures 3 and 4).

**Table 8.**
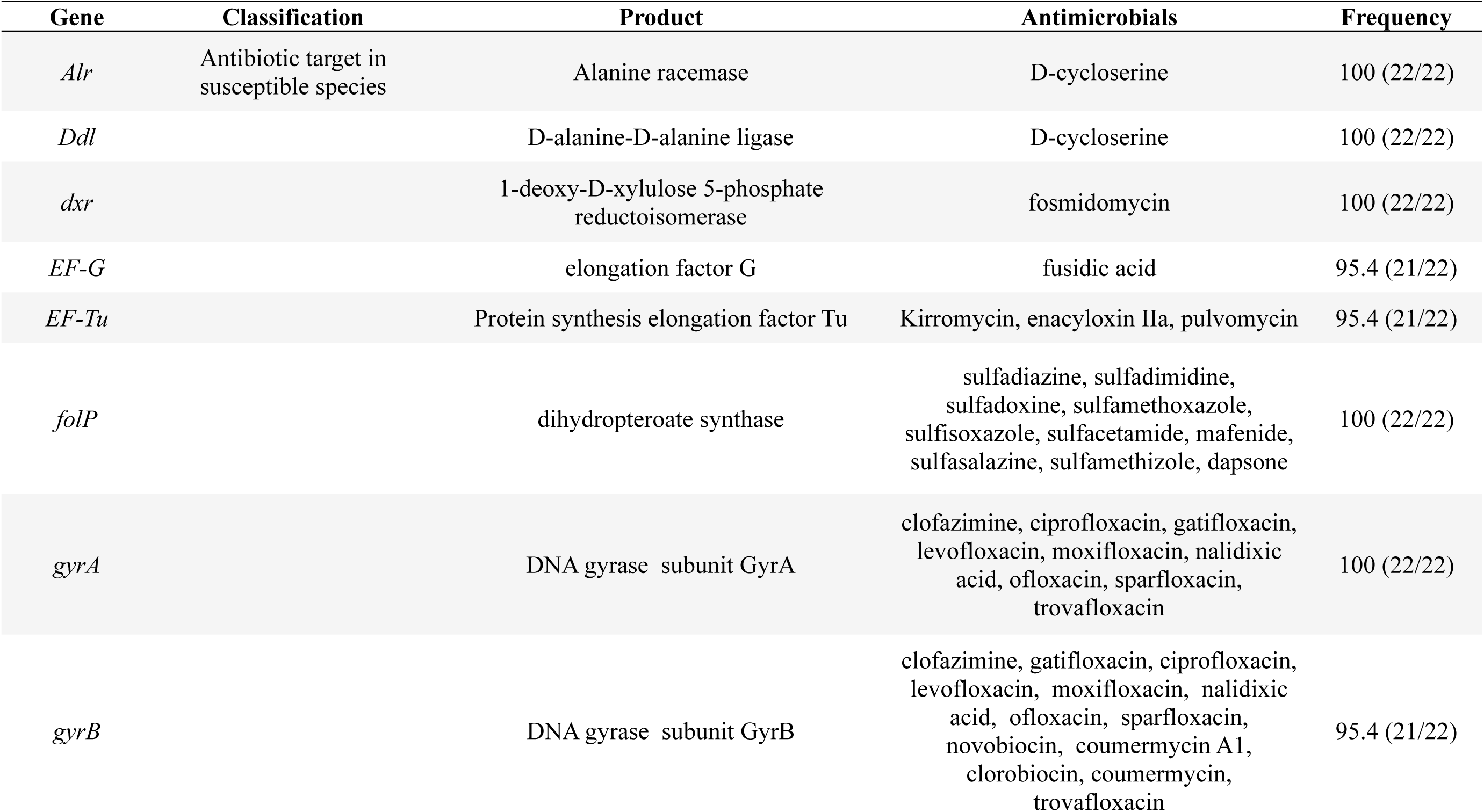
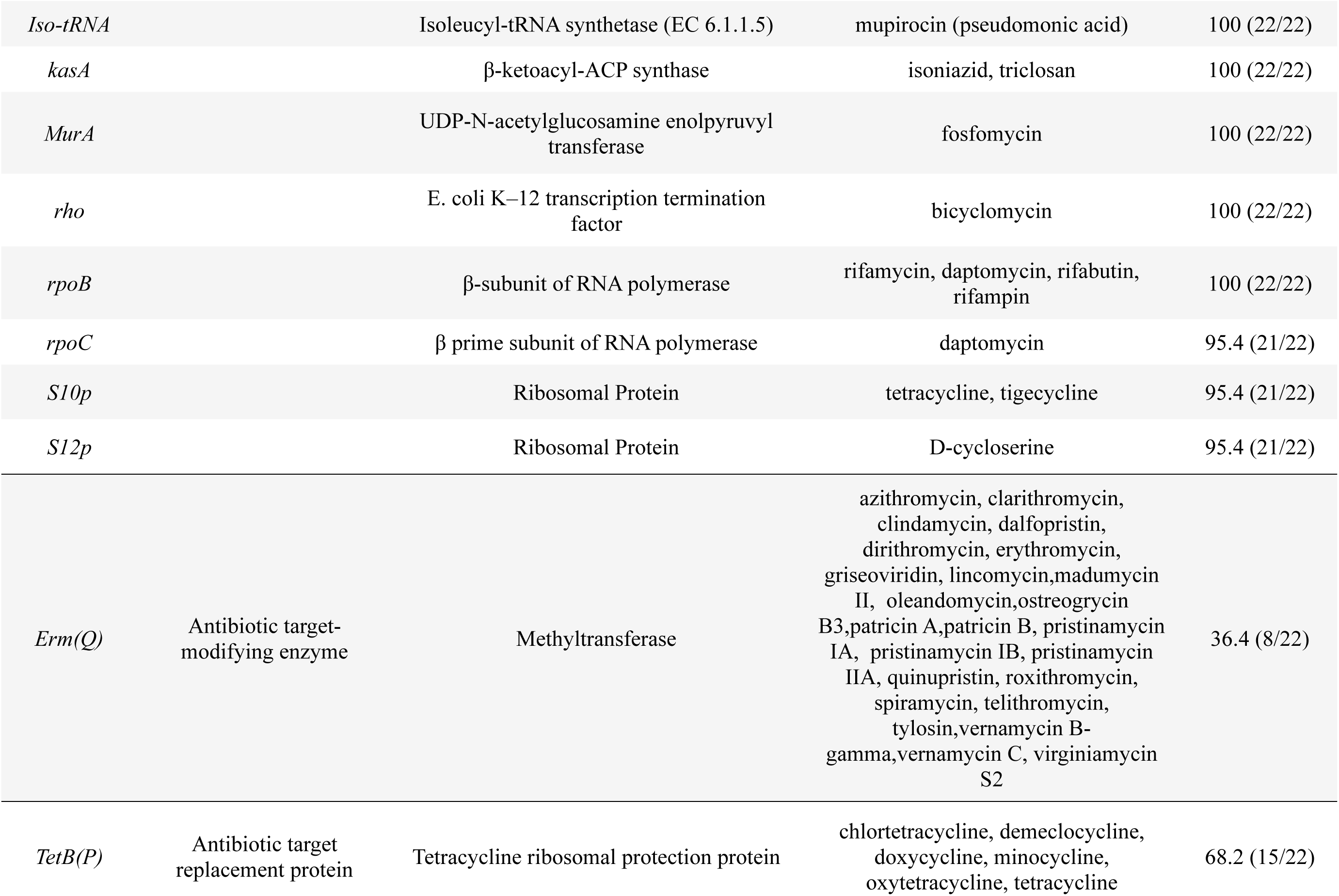
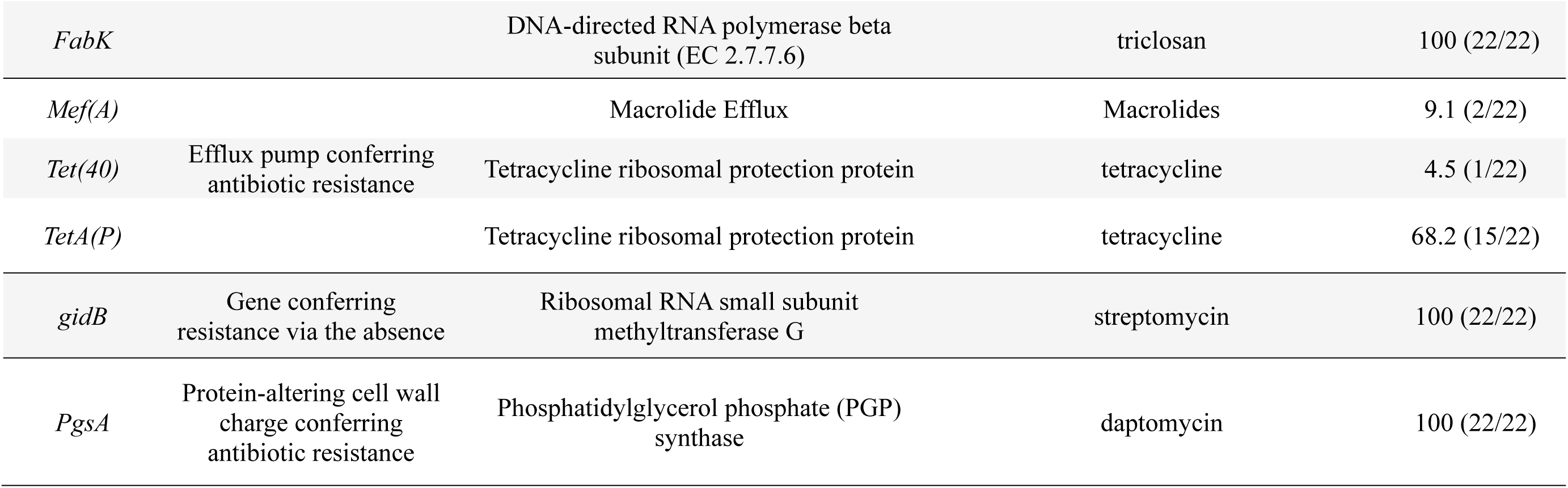
Antibiotic Target Genes Identified in *Peptacetobacter hiranonis* strains. This table summarizes the antibiotic target genes detected in *Peptacetobacter hiranonis* strains. It includes the gene name, classification, gene product, associated antimicrobials, and the frequency of detection across the strains analyzed.\

Description of Peptacetobacter hiranonis subsp. deconjugans subsp. nov.

*Peptacetobacter hiranonis* subsp. *deconjugans* (de.con.ju’ga.ns. N.L. participle adjective deconjugans, deconjugating, referring to the strains’ ability to deconjugate bile acids). The bile acid-converting bacterium *P. hiranonis* subsp. *deconjugans* are commonly isolated from the feces of healthy adult dogs and present the choloylglycine hydrolase activity, which catalyzes the deconjugation of glycine- and taurine-conjugated bile acids. This description is based on 12 strains. Cells are Gram-stain-positive, non-motile, potentially spore-forming, straight, and rod-shaped. Grows at 37°C on Brucella Blood and Brain-Heart Infusion (BHI) agar and Brucella, Brain-Heart Infusion, Peptone-Yeast Glucose, and Chopped Meat broth. On Brucella blood agar, colonies were grey to off-white in color, 1-3 mm in diameter, slightly convex, with irregular edges, and non-hemolytic after 24 hours of anaerobic incubation at 37°C. Distinguishes from *P. hiranonis* subsp. *nondeconjugans* for its ability to deconjugate bile acids. Utilized glucose, saccharose, and mannose. Lactose, maltose, salicin, xylose, arabinose, glycerol, cellobiose, esculin, melibiose, raffinose, rhamnose, trehalose, sorbitol, gelatin hydrolysis, indole production, urease, catalase, or hydrogen sulfide (H₂S) negative. Variable results for mannitol. Positive for N-Acetyl- ßglucosaminidase. Negative for α-galactosidase, ß-galactosidase, ß-galactosidase 6-phosphatase, α-glucosidase, ß-glucosidase, α-arabinosidase, ß-glucuronidase, α-fucosidase, phenylalanine arylamidase, tyrosine arylamidase, glycine arylamidase, histidine arylamidase, glutamyl-glutamic acid arylamidase, and serine arylamidase. The type strain, *Peptacetobacter hiranonis* subsp. *deconjugans* CH8, is held in the Agricultural Research Culture Collection (NRRL), and the accession number is being processed. The genome of the type strain is associated with BioProject accession number PRJNA1516050 in GenBank/EMBL.

### Description of Peptacetobacter hiranonis subsp. nondeconjugans subsp. nov

*Peptacetobacter hiranonis* subsp*. nondeconjugans* (non.de.con.ju’ga.ns. N.L. participle adjective nondeconjugans, not deconjugating; referring to the strains’ inability to deconjugate bile acids). The bile acid-converting bacterium *P. hiranonis* subsp. *nondeconjugans* are commonly isolated from the feces of healthy adult cats and do not present the choloylglycine hydrolase activity, which catalyzes the deconjugation of glycine- and taurine-conjugated bile acids. The description is based on 10 strains. Cells are Gram-stain-positive, non-motile, potentially spore-forming, straight, and rod-shaped. Grows at 37°C on Brucella Blood and Brain-Heart Infusion (BHI) agar and Brucella, Brain-Heart Infusion, Peptone-Yeast Glucose, and Chopped Meat broth. On Brucella blood agar, colonies were grey to off-white in color, 1-3 mm in diameter, slightly convex, with irregular edges, and non-hemolytic after 24 hours of anaerobic incubation at 37°C. Utilized glucose, saccharose, mannitol, and mannose. Lactose, maltose, salicin, xylose, arabinose, glycerol, cellobiose, esculin, melibiose, raffinose, rhamnose, trehalose, gelatin hydrolysis, indole production, urease, catalase, or hydrogen sulfide (H₂S) negative. Variable results for sorbitol. Positive for N-Acetyl- ßglucosaminidase, arginine, and alanine arylamidase. Negative for α-galactosidase, ß- galactosidase, ß-galactosidase 6-phosphatase, α-glucosidase, ß-glucosidase, α-arabinosidase, ß- glucuronidase, α-fucosidase, phenylalanine arylamidase, tyrosine arylamidase, glycine arylamidase, histidine arylamidase, glutamyl-glutamic acid arylamidase, and serine arylamidase. The type strain, *Peptacetobacter hiranonis* subsp. *deconjugans* CH17, is held in the Agricultural Research Culture Collection (NRRL), and the accession number is being processed. The genome of the type strain is associated with BioProject accession number PRJNA1516050 in GenBank/EMBL.

## Author Notes

The genomes of *Peptacetobacter hiranonis* strains are associated with BioProject accession number PRJNA1516050 in GenBank/EMBL. The supplementary figures, tables, and supplementary files are available with the online version of this article.

## Abbreviations

BA: Bile Acid
BSH: Bile Salt Hydrolase
bai: Bile Acid-Inducible
HSDH: Hydroxysteroid Dehydrogenase
CA: Cholic Acid
CDCA: Chenodeoxycholic Acid
DCA: Deoxycholic Acid
LCA: Lithocholic Acid
dDDH: Digital DNA-DNA Hybridization
ANI: Average Nucleotide Identity
MIC: Minimum Inhibitory Concentration

## Ethical Statement

The Institutional Animal Care and Use Committee (IACUC) at Texas A&M University Division of Research has determined that receiving fecal samples passively collected by owners of privately-owned animals, as no animals will be handled or disturbed as a function of research, no Animal Use Protocol was required.

## Author Contributions

Bruna Correa Lopes: study design, data collection, data analysis, and writing – initial draft and review of final version. Jonathan Turck: data collection, data analysis, and writing – review of final version. Amanda B. Blake: data analysis and writing – review of final version. Luis Fernando da Costa Medina: data collection and writing – review of final version. Sara Lawhon: study design and writing – review of final version. Jan S. Suchodolski: study design, funding acquisition, and writing – review of final version. And Rachel Pilla: study design, funding acquisition, and writing – review of final version.

## Acknowledgments

We appreciate the collaboration with Dr. Joseph Sorg for providing the *Clostridium scindens* wild-type strain. Portions of this research were conducted with the advanced computing resources provided by Texas A&M High Performance Research Computing.

## Data Availability Statement

Additional data supporting the findings of this study are available from the corresponding author upon reasonable request.

## Supplementary Material

Supplementary File 1. Composition of Brucella (BRU) Blood Agar by Anaerobe Systems (California, USA).

Supplementary Figure 1. Scatter plots representing the digital DNA-DNA hybridization (dDDH) percentages for *Peptacetobacter hiranonis* subsp. *deconjugans* (circles) and *P. hiranonis* subsp. *nondeconjugans* (triangles) isolated from cats (left) and dogs (right), calculated using three different dDDH formulae (d0, d4, and d6) in comparison with the reference strain *P. hiranonis* GCF_016694795.1. Each panel represents results obtained with a specific dDDH formula. Red horizontal bars indicate median values. The shaded area denotes the 70–100% dDDH threshold range typically considered indicative of the same species.

Supplementary Figure 2. Alignment lengths and Hadamard matrix for *Peptacetobacter hiranonis* ANIm analysis.

Supplementary Figure 3. Frequency distribution for Minimum Inhibitory Concentration (MIC) for sixteen antimicrobials (named: amoxicillin, amoxicillin/clavulanate, cephalothin, chloramphenicol, metronidazole, streptomycin, azithromycin, cefepime, ciprofloxacin, clindamycin, tetracycline, vancomycin, cefoxitin, ceftriaxone, daptomycin, and meropenem) tested against *Peptacetobacter hiranonis* strains.

Supplementary Figure 4. Frequency of strains with Minimum Inhibitory Concentration (MIC) ≤ or > 2 µl/mL based on the presence or absence of antimicrobial resistance (AMR) genes detected on the genome of the *Peptacetobacter hiranonis* strains.

Supplementary Table 1. Identification, year, source of isolation, host species, region of residence, breed, sex, and age of animals from which samples were obtained for all strains of *Peptacetobacter hiranonis* included in this study.

Supplementary Table 2. CheckM results for the assembled genomes of *Peptacetobacter hiranonis*.

Supplementary Table 3. Comprehensive Genome Analysis (CGA) by BV-BRC of the assembled genome of *Peptacetobacter hiranonis* strains.

Supplementary Table 4. Genome-to-Genome Distance Comparison Between *Peptacetobacter hiranonis* subsp. *deconjugans* and subsp. *nondeconjugans* Strains. This table presents the results of digital DNA–DNA hybridization (dDDH) analyses between *Peptacetobacter hiranonis* subsp. *deconjugans* and subsp. *nondeconjugans* query strains and reference genome, P. hiranonis (GCF_016694795.1). Three dDDH formulas (d0, d4, and d6) and their corresponding confidence intervals (C.I.) are shown, along with the difference in G+C content between genomes.

Supplementary Table 5. Biochemical profile of *Peptacetobacter hiranonis* subsp. *deconjugans* and subsp. *nondeconjugans* assessed using API20A and API32 A Rapid (BioMérieux).

Supplementary Table 6. Median, minimum, and maximum values for the Minimum Inhibitory Concentration (MIC) of *Peptacetobacter hiranonis* subsp. *deconjugans* and subsp. *nondeconjugans.* MIC was assessed using ETEST^®^.

Supplementary Table 7. Mean values for bile acid deconjugation and conversion, and presence and absence of bai operon genes in *Peptacetobacter hiranonis* subsp. *deconjugans* and subsp. *nondeconjugans*.

