## Supplementary_material for "Description of canine- and feline-derived strains of the bile acid-converting bacterium *Peptacetobacter hiranonis: P. hiranonis* subsp. *deconjugans* subsp. nov. and *P. hiranonis* subsp. *nondeconjugans* subsp. nov.": Supplementary_Figures_BCL_07142026.pdf

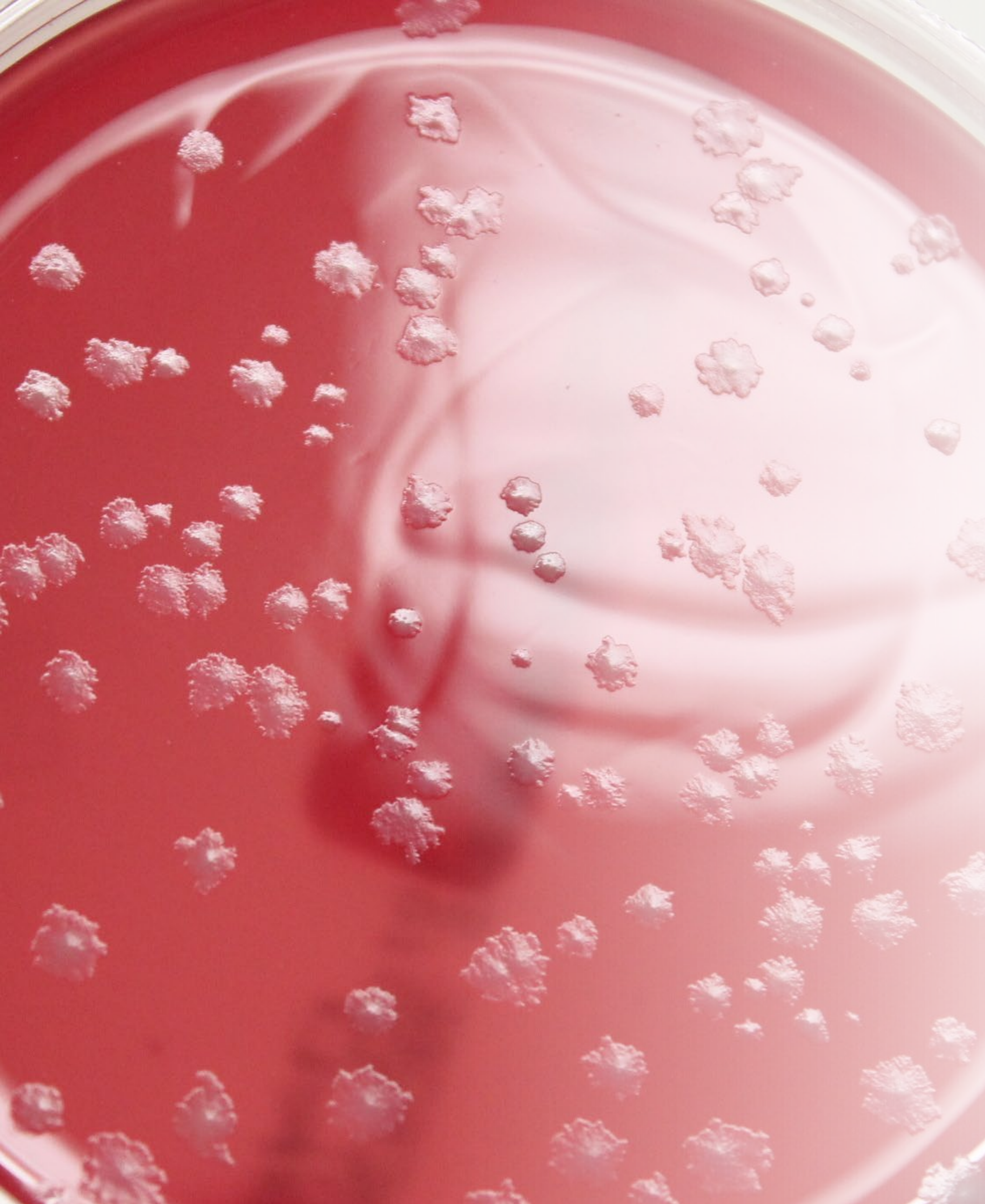

### Description of canine- and feline-derived strains of the bile acid-converting bacterium *Peptacetobacter hiranonis: P. hiranonis* subsp. *deconjugans* subsp. nov. and *P. hiranonis* subsp. *nondeconjugans* subsp. nov.

Bruna Correa Lopes[1], Jonathan Turck[1], Amanda Blake[1], Luis Fernando da Costa Medina[1], Sara D. Lawhon[2], Jan S. Suchodolski[1], and Rachel Pilla[3]

[1]Gastrointestinal Laboratory, Department of Small Animal Clinical Sciences, Texas A&M University, College Station, TX, USA

[2]Departments of Veterinary Pathobiology, College of Veterinary Medicine and Biomedical Sciences, Texas A&M University, College Station, TX, USA

[3]Department of Veterinary Medicine and Animal Sciences, University of Milan, Italy

**Supplementary Figure 1.** Scatter plots representing the digital DNA-DNA hybridization (dDDH) percentages for *Peptacetobacter hiranonis* subsp. deconjugans (circles) and *P. hiranonis* subsp. nondeconjugans (triangles) isolated from cats (left) and dogs (right), calculated using three different dDDH formulae (d0, d4, and d6) in comparison with the reference strain *P. hiranonis* GCF_016694795.1. Each panel represents results obtained with a specific dDDH formula. Red horizontal bars indicate median values. The shaded area denotes the 70–100% dDDH threshold range typically considered indicative of the same species.

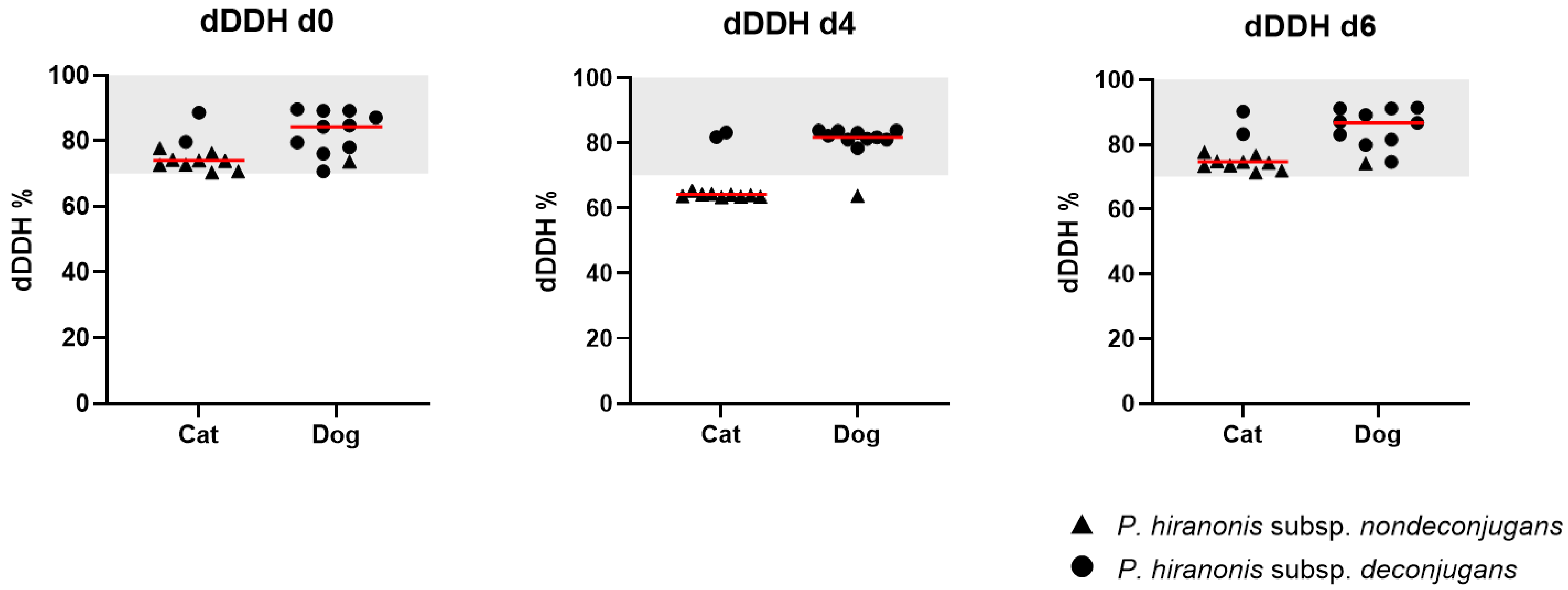

**Supplementary Figure 2.** Alignment lengths (a) and Hadamard (b) matrix for *Peptacetobacter hiranonis* ANIm analysis.

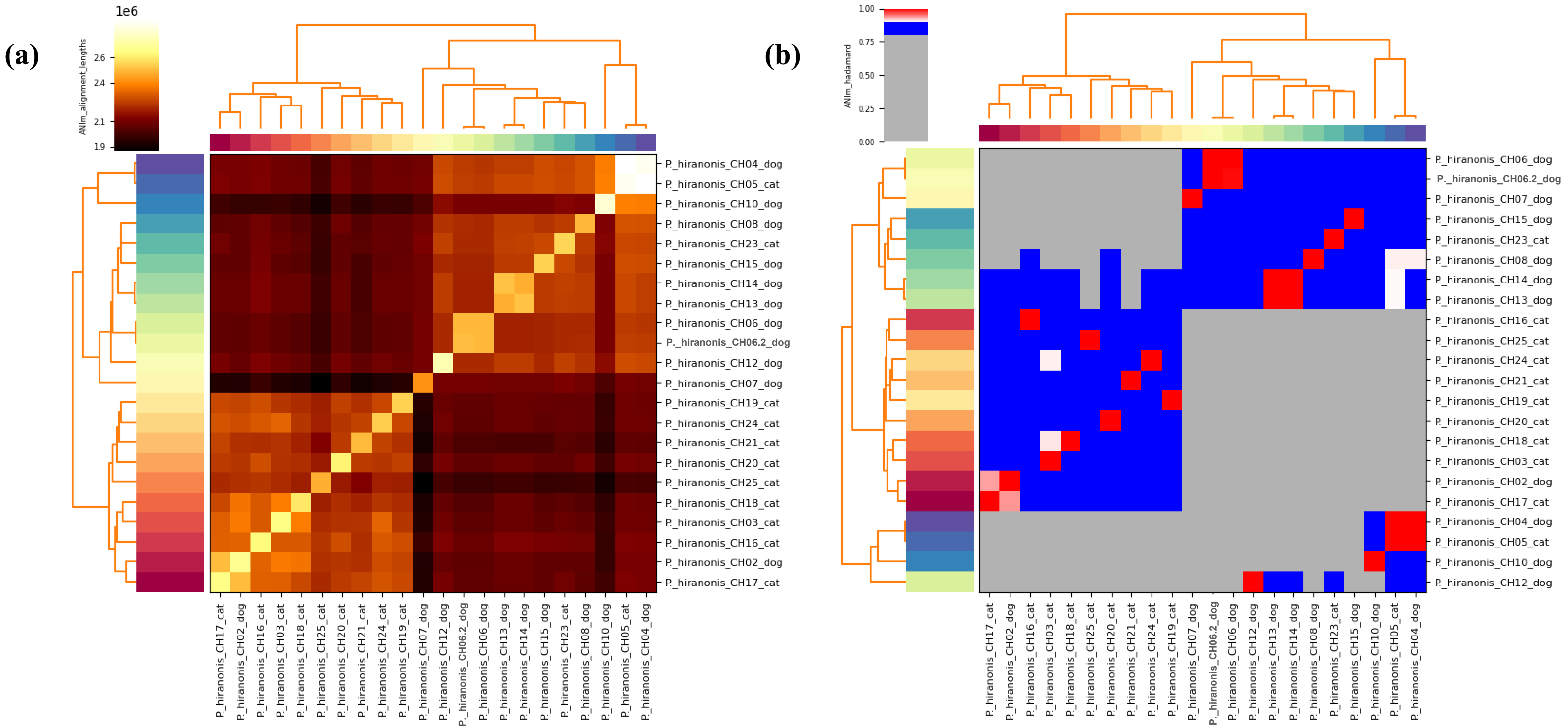

**Supplementary Figure 3.** Frequency distribution for Minimum Inhibitory Concentration (MIC) for sixteen antimicrobials (named: amoxicillin, amoxicillin/clavulanate, cephalothin, chloramphenicol, metronidazole, streptomycin, azithromycin, cefepime, ciprofloxacin, clindamycin, tetracycline, vancomycin, cefoxitin, ceftriaxone, daptomycin, and meropenem) tested against *Peptacetobacter hiranonis* strains.

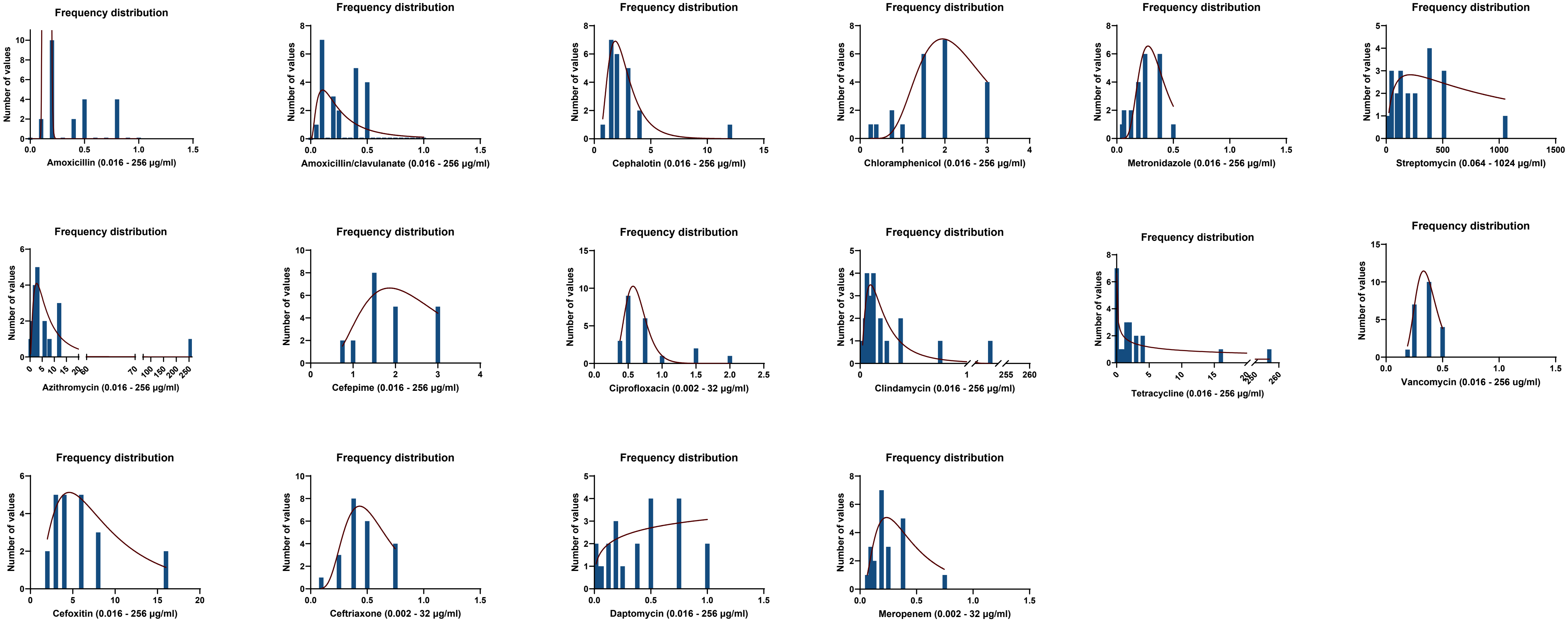

**Supplementary Figure 4.** Frequency of strains with Minimum Inhibitory Concentration (MIC) ≤ or > 2 µl/mL based on the presence or absence of antimicrobial resistance (AMR) genes detected on the genome of the *Peptacetobacter hiranonis* strains.

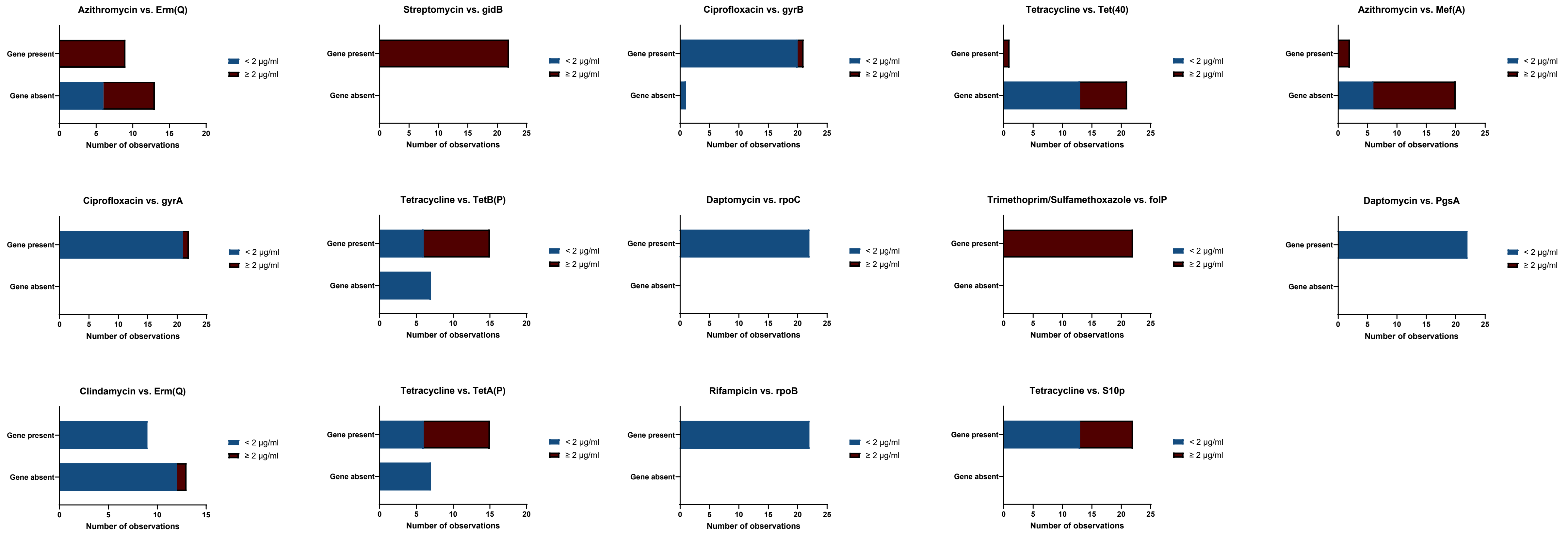
