## Supplementary_material for "Description of canine- and feline-derived strains of the bile acid-converting bacterium *Peptacetobacter hiranonis: P. hiranonis* subsp. *deconjugans* subsp. nov. and *P. hiranonis* subsp. *nondeconjugans* subsp. nov.": Supplementary_file_BCL_04192026.pdf

**Supplementary File 1.** Composition of Brucella (BRU) Blood Agar by Anaerobe Systems (California, USA).

**Summary:**

BRU agar is an enriched non-selective medium that supports the growth of fastidious microorganisms. BRU agar contains casein, peptones, yeast extract, and dextrose as the nutrient base medium. It is supplemented with vitamin K1 and hemin to facilitate the recovery of fastidious anaerobes. Sheep blood has been added for the growth factors required by some anaerobic bacteria, and allows the observation of hemolytic reactions as seen by the double zone β-hemolysis of *Clostridium perfringens*, for example. This media is prepared, dispensed, and packaged under oxygen-free conditions to prevent the formation of oxidized products prior to use.

**Formulation:**

| | |
|---|---|
| Pancreatic Digest of Casein | 10 g |
| Soy Peptone | 3 g |
| Meat Peptone | 10 g |
| Dextrose | 1 g |
| Yeast Extract | 2 g |
| Sodium Chloride | 5 g |
| Sodium Bisulfite | 0.1 g |
| L-Tryptophan | 0.2 g |
| Calcium Lactate | 0.5 g |
| Sodium Acetate | 0.5 g |

| | |
|---|---|
| Ascorbic Acid | 0.1 g |
| Hemin (0.1% solution) | 5 mL |
| Vitamin K1 (1.0% solution) | 1 mL |
| L-Cystine | 0.4 g |
| Sodium Hydroxide (4.0% solution) | 4 mL |
| Agar | 15 g |
| Sheep Blood | 45.5 mL |
| DI Water | 1 L |

*Approximate formula. Adjusted and/or supplemented as required to meet performance criteria.

Final pH: 7.1 ± 0.4 at 25° C

Final weight: 16.0 g ± 1.6 g
